# Comparative RNA profiling identifies stage-specific phasiRNAs and co-expressed *Argonaute* genes in Bambusoideae and Pooideae species

**DOI:** 10.1101/2024.06.12.598608

**Authors:** Sébastien Bélanger, Junpeng Zhan, Yunqing Yu, Blake C. Meyers

## Abstract

PhasiRNAs (phased, small interfering RNAs) play a crucial role in supporting male fertility in grasses. Earlier work in maize (*Zea mays*) and rice (*Oryza sativa*) – and subsequently many other plant species – identified premeiotic 21-nt and meiotic 24-nt phasiRNAs. More recently, a group of premeiotic 24-nt phasiRNAs were discovered in the anthers of two Pooideae species, barley (*Hordeum vulgare*) and bread wheat (*Triticum aestivum*). The conservation of premeiotic 24-nt phasiRNAs and other classes of reproductive phasiRNAs across Pooideae species remains unclear. We conducted a comparative RNA profiling of three anther stages in six Pooideae species and one Bambusoideae species. We observed complex temporal accumulation patterns of 21-nt and 24-nt phasiRNAs in Pooideae and Bambusoideae grasses. In Bambusoideae, 21-nt phasiRNAs accumulated during meiosis, whereas 24-nt phasiRNAs were present in both premeiotic and postmeiotic stages. We identified premeiotic 24-nt phasiRNAs in all seven species examined. These phasiRNAs exhibit distinct biogenesis mechanisms and potential Argonaute effectors compared to meiotic 24-nt phasiRNAs. We show that specific *Argonaute* genes co-expressed with stage-specific phasiRNAs are conserved across Bambusoideae and Pooideae species. Our degradome analysis identified a set of conserved miRNA target genes across species, while 21-nt phasiRNAs targets were species-specific. Cleavage of few targets was observed for 24-nt phasiRNAs.

## INTRODUCTION

Plants produce small RNAs (sRNAs) of 20 to 24 nucleotides (nt) in length, categorized based on their length, biogenesis, or functions. These sRNAs can be grouped into two main categories: microRNAs (miRNAs) and small interfering RNAs (siRNAs). miRNAs originate from a single-stranded transcript synthesized by polymerase II (Pol II) which forms stem-loop structures that are cleaved by a ribonuclease III enzyme called Dicer-like 1 (DCL1) to produce miRNAs, which are 20-22 nt in length (Axtell and Meyers 2018). miRNAs play a crucial role in the regulation of various biological processes, primarily through post-transcriptional gene regulation of target mRNAs and long non-coding RNA transcripts (Borges and Martienssen 2015; Axtell and Meyers 2018). Generally, cleavage of protein-coding or long non-coding RNAs (lncRNAs) by 21-nt miRNAs leads to target degradation while cleavage by 22-nt miRNAs triggers the production of 21-nt or 24-nt phased secondary siRNAs (phasiRNAs) (Liu et al. 2020).

Previous work in rice (*Oryza sativa*; Oryzoideae) and maize (*Zea mays*; Panicoideae) classified phasiRNAs into two groups of 21-nucleotide (nt) and 24-nt phasiRNAs. While the 21-nt phasiRNAs participate in both vegetative and reproductive development, so far, 24-nt phasiRNAs have been exclusively observed in male reproductive development (Teng et al. 2020). In rice, the biogenesis of reproductive 21-nt phasiRNAs typically initiates by miR2118-directed Argonaute 1d (AGO1d)-catalyzed cleavage of phasiRNA precursor (*PHAS* precursors) transcripts that are converted into double-stranded RNA (dsRNA) by RNA-directed RNA polymerase 6 (RDR6), processed into 21-nt sRNA duplexes by DCL4, and loaded onto Argonaute effectors (AGO1d or AGO5c) to target specific genes in a posttranscriptional gene silencing (PTGS) manner (Song et al. 2012a, 2012b; Komiya et al. 2014; Shi et al. 2022). The 24-nt phasiRNA pathway in grasses relies on AGO1d, miR2275, RDR6 and DCL5 (Song et al. 2012a, 2012b; Komiya et al. 2014; Teng et al. 2020; Shi et al. 2022), but their AGO effectors and downstream regulatory mechanisms remain largely unknown.

More recently, our group has described a group of premeiotic 24-nt phasiRNAs that accumulate in anthers of barley (*Hordeum vulgare*) and bread wheat (*Triticum aestivum*), both belonging to the Triticeae tribe (Pooideae) (Figure 1A), and then in maize, teosinte and rice (Bélanger et al. 2020; Zhan et al. 2024). The number of loci encoding premeiotic 24-nt phasiRNAs in barley and wheat is significantly higher than in maize, teosinte and rice, suggesting their expansion in the Triticeae (Bélanger et al. 2020; Zhan et al. 2024). In barley and wheat, a conserved motif shared among thousands of premeiotic 24-*PHAS* precursors does not match with the miR2275 motif conserved in meiotic *PHAS* precursors, suggesting that another sRNA mediates the biogenesis of the premeiotic group (Bélanger et al. 2020). The conserved motif identified among tens of premeiotic 24-*PHAS* precursors in maize and teosinte differs from the motif described in barley and wheat (Zhan et al. 2024). In maize and rice, meiotic 24-*PHAS* precursors are expressed in the tapetum (a specialized layer of nutritive cells found within the anther) while 24-nt phasiRNAs accumulate in the tapetum and meiocytes (developing pollen grains) (Zhai et al. 2015; Fei et al. 2016). However, premeiotic 24-nt phasiRNAs peak before the differentiation of the tapetum and meiotic cells. A gene coexpression analysis describing the regulation of phasiRNA biogenesis components in wheat and barley anthers highlighted AGO4c (aka AGO9) and AGO6 as candidate binding partners of premeiotic and meiotic 24-nt phasiRNAs, respectively (Bélanger et al. 2020). Thus, our previous work provides multiple lines of evidence that premeiotic 24-*PHAS* loci have a distinct evolutionary trajectory, stage-specific biogenesis and spatial expression compared to the meiotic group.

**Figure 1:**
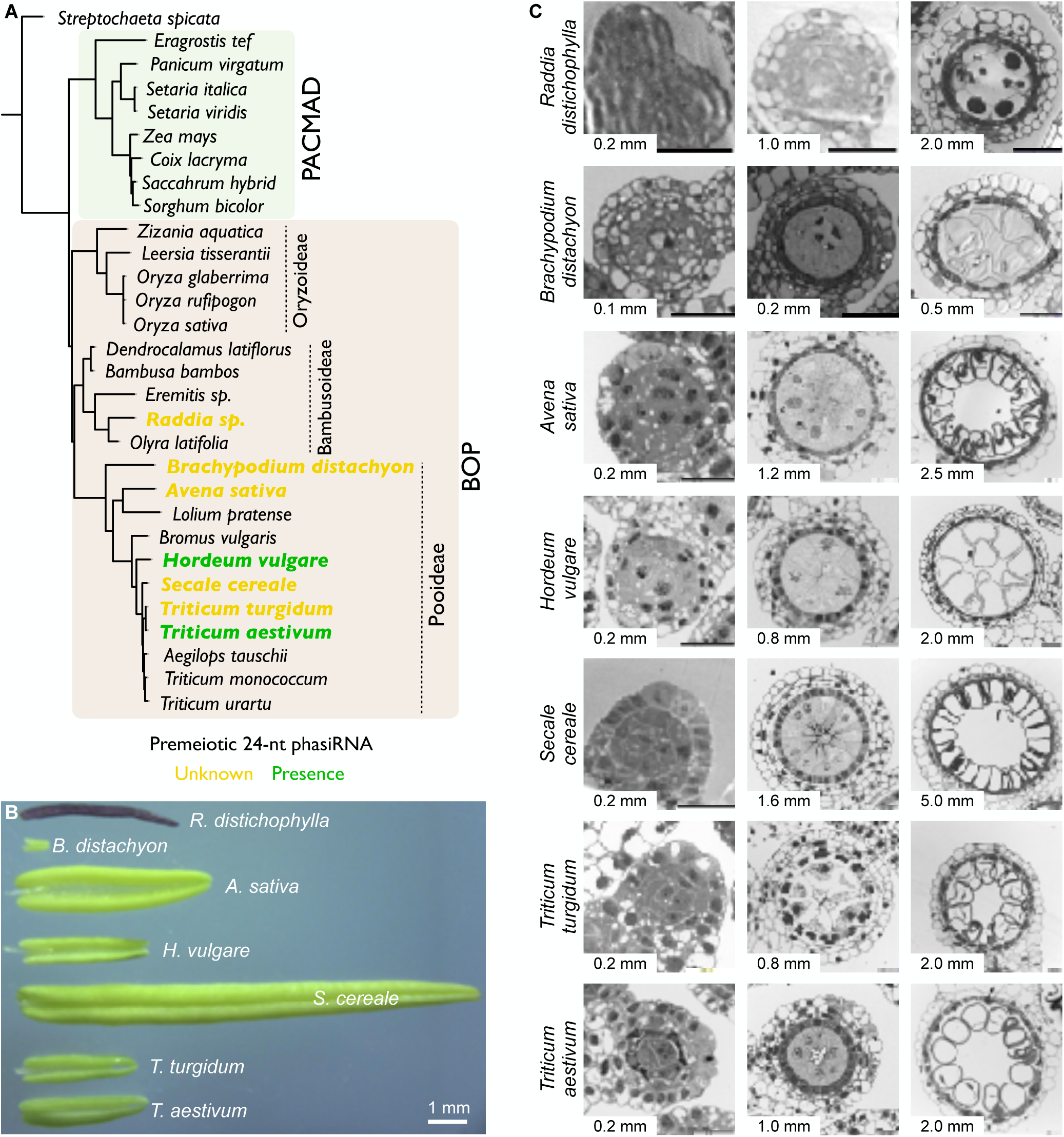
Species and anther stages sampled for RNA experiments. (A) Phylogenetic tree representing the evolutionary relationship of a subset of Poaceae species, modified from Saarela *et al*. (Saarela et al. 2018). Previous publications from our group have described the presence (in green) of pre-meiotic 24-nt phasiRNAs. To further characterize premeiotic 24-nt phasiRNAs, we sampled two species known to express these phasiRNAs (in green) and five species with an unknown status (in yellow). (B) Anthers at the late stage of pollen maturation, before anthesis, exhibit significant variation in length. The shortest anther measures 0.7 mm (in *B. distachyon*), while the longest anther measures 12.5 mm (in *S. cereale*). (C) Anthers sampled from one Bambusoideae (*R. distichophylla*) and six Pooideae (*B. distachyon*, *A. sativa*, *H. vulgare*, *S. cereale*, *T. turgidum*, and *T. aestivum*) species. The details of each sampled variety are provided in Supplemental Table 1. Anthers were fixed with a 2% (v/v) paraformaldehyde and 2% (v/v) glutaraldehyde solution, embedded using the Quetol epoxy resin, sectioned to 500 µm, and stained using the Epoxy Tissue Stain solution. Scale bar: 20 µm.

To date, the conservation of premeiotic 24-nt phasiRNAs and other classes of reproductive phasiRNAs across Pooideae species, and even across angiosperms, remains unclear. Moreover, there is limited knowledge about the regulatory mechanism and function of 21-nt phasiRNA and groups of premeiotic and meiotic of 24-nt phasiRNAs; it is still unclear if 24-nt phasiRNAs recognize specific RNA transcripts to guide AGO-catalyzed cleavage of transcripts. Therefore, we conducted transcriptome and degradome analyses to investigate the evolution, conservation, regulation, and function of 21-nt and 24-nt phasiRNAs in the Pooideae subfamily. We collected samples from anthers at premeiotic, meiotic, and postmeiotic stages of seven species, including six Pooideae species and one Bambusoideae as an outgroup (Figure 1A and Supplemental Table 1). Our study revealed distinct temporal accumulation of 21-nt and 24-nt phasiRNAs in anthers between Pooideae and Bambusoideae species. We found that while 21-nt phasiRNAs are exclusively or dominantly present in the premeiotic anthers of all six Pooideae species, they are mostly present at the meiotic stage in *Raddia distichophylla* (Bambusoideae). Moreover, our data revealed the presence of 24-nt phasiRNAs in the premeiotic anthers of all seven species and these are distinct from the meiotic 24-nt phasiRNAs in their biogenesis mechanisms and functions. Finally, the degradome analysis demonstrates that miRNA targets are quite conserved across species, targets of 21-nt phasiRNAs are species-specific, and nearly no targets could be validated for 24-nt phasiRNAs.

## RESULTS

### Diverse patterns of anther developmental progression in the Pooideae and Bambusoideae lineages

To investigate premeiotic 24-nt phasiRNAs and other classes of reproductive phasiRNAs, we collected anthers from one Bambusoideae (*R. distichophylla*) and six Pooideae (*Brachypodium distachyon*, *Avena sativa*, *H. vulgare*, *Secale cereale*, *T. turgidum*, and *T. aestivum*) species to examine the accumulation of phasiRNAs during premeiotic, meiotic, and postmeiotic stages of anther development (Figure 1A and Supplemental Table 1). The length of the anther serves as an effective marker to determine the premeiotic, meiotic, and postmeiotic stages in a given grass species and variety. Therefore, we imaged cross-sections of anthers (as schematized in Supplemental Figure 1A) to examine the developmental progression of meiosis at 7 time points for *Brachypodium*, 13 time points for barley, herbaceous bamboo (*R. distichophylla*), durum wheat (*T. turgidum*), and bread wheat, 15 time points for oat (*A. sativa*), and 18 time points for rye (*S. cereale*).

We observed a significant disparity in anther length at anthesis, with the shortest 0.7 mm in *Brachypodium* and the longest measuring 12.5 mm in rye (Figure 1B). Among Pooideae species, except for *Brachypodium*, we observed no notable differences during the early stages of anther development. Cell fate specification was observed in anthers measuring 0.2 mm and 0.4 mm, followed by the emergence of four distinct cell layers, namely the epidermis, endothecium, middle layer, and tapetum, by the 0.6 mm stage (Supplemental Figure 1). At 0.6 mm, microspore mother cells developed from sporogenous tissues, and meiosis I initiation occurred at the 0.8 mm stage in all species except *Brachypodium*. The tapetum reached its maximum size in terms of cell width during the early stage of meiosis I, and then gradually degraded after meiosis during the pollen maturation stage. In contrast to the premeiotic stages, the progression of anther development during meiosis I and II varied among barley, wheat (durum and bread), oat, and rye species. Indeed, the stage of uninuclear vacuolate microspores (early postmeiotic) was observed at 1.8 mm (barley and wheat species), 2.5 mm (oat), and 5.0 to 7.5 mm (rye) stages (Figure 1C and Supplemental Figure 1 C–G). Subsequently, we observed a slower developmental process characterized by an extended length increment during pollen maturation. Vacuolated microspores matured into pollen grains, which were released upon dehiscence at stages of 3.0 to 3.5 mm (barley and wheat species), 5.0 mm (oat), or 12.5 mm (rye) (Supplemental Figure 1 C–G). In the herbaceous bamboo, we harvested premeiotic, meiotic, and postmeiotic stages at 0.2 mm, 1.0 mm, and 2.0 mm, respectively, while pollen was shed at 4.5 mm (Figure 1C). Notably, the anthers of the bamboo species were pigmented purple, unlike the green shade observed in other species (Figure 1B). Anther development in *Brachypodium* was remarkably rapid, as the premeiotic (≤0.1 mm), meiotic (0.2 mm to 0.4 mm), and postmeiotic (0.5 mm to 0.7 mm) stages were completed at 0.7 mm anthers. Based on the aforementioned observations, we sequenced 189 RNA libraries (evenly distributed across sRNAs, transcripts, and uncapped transcripts), totaling 4.03 billion reads from anthers at premeiotic, meiotic, and postmeiotic stages specific to each species (Figure 1C).

### Numerous loci express miRNAs but represent a small number of MIR families in anthers

Prior to analyzing phasiRNAs, we used the sRNA libraries to annotate miRNA and their abundance levels in developing anthers, comparing the overlap in miRNAs expressed among species. We identified from 54 to 164 miRNA loci in the Pooideae species and a total of 58 loci in bamboo. Supplemental Table 2 summarizes the number of miRBase-annotated miRNAs and previously unannotated miRNAs among sampled species. Supplemental Table 3 details the coordinates and abundances of miRNAs annotated from anther tissues. Known miRNAs represented a relatively small number of *MIR* families per species (16 to 27 families). Notably, only 13 families were shared among all species (Supplemental Figure 2) and these are known as broadly conserved across the land plants (e.g. miR156, miR166/167/168, miR169, miR171, miR172, miR319, miR390, miR393, miR396). Both miR2118 and miR2275 families, which mediate cleavage of 21-*PHAS* and 24-*PHAS* precursor transcripts, respectively, are expressed in anthers of all species (Supplemental Figure 2). The number of MIR2118 and MIR2275 copies vary substantially among the sampled species, with the number of MIR2118 copies ranging from eight (in *Brachypodium* species) to 28 (in durum wheat species), and MIR2275 ranging from two (in *Brachypodium*) to 12 (in oat and bread wheat). Notably, we observed a total of nine and four *MIR* families found only in *Brachypodium* and bamboo anthers, respectively (Supplemental Figure 2). One to three uniquely expressed *MIR* families were found in other species except oat (Supplemental Figure 2). We designated miRNAs that could not be annotated with miRBase (Kozomara and Griffiths-Jones 2014; Kozomara et al. 2018) as “candidate” (Supplemental Table 2 and 3). The majority of these candidate miRNAs appear to be newly evolved, genus- or species-specific miRNAs, while the majority of miRNAs that are annotated by miRBase were shared in two or more species (except for *Brachypodium* and bamboo).

### The number of PHAS loci increases with the size and ploidy of genomes

We identified from 1,242 to 13,541 *PHAS* loci in the Pooideae species and annotated a total of 1,019 in bamboo (Supplemental Table 4 and detailed in 5). The number of *PHAS* loci increased with the genome size in diploid species and with ploidy in the polyploids. Diploid genomes generated phasiRNAs from 1,019 1,242, 2,634 and 7,672 *PHAS* loci in bamboo, *Brachypodium*, barley, and rye genomes, respectively. Similarly, the tetraploid durum wheat genome had 8,553 *PHAS* loci while the hexaploid oat and bread wheat genomes generated phasiRNAs from 13,452 and 13,541 *PHAS* loci, respectively (Supplemental Table 4). *PHAS* loci annotated in polyploid genomes exhibited similar distribution patterns across homoeologous chromosomes. The ratio of 21-*PHAS* and 24-*PHAS* loci differs between genomes. In oat, barley and wheat genomes, there were 2-fold to 3-fold more 21-*PHAS* than 24-*PHAS* loci and this ratio was consistent between the oat and wheat subgenomes. That ratio was higher in the bamboo, *Brachypodium* and rye genomes in which we observed as much as 3.9-fold more 21-*PHAS* than 24-*PHAS* loci in bamboo (Supplemental Table 4). Together, these results indicate that the number of *PHAS* loci increases with the size and ploidy of these grass genomes.

### Temporal accumulation of 21-nt phasiRNAs in anthers of Pooideae

Our analysis on 21-nt phasiRNAs revealed their distinct accumulation patterns in bamboo compared to Pooideae species studied here (Figure 2A), and previously described in maize and rice (Zhai et al. 2015; Fei et al. 2016). In bamboo, most *PHAS* loci generate 21-nt phasiRNAs with peak abundance at the meiotic stage of anthers (96.7%), while in the Pooideae, the majority of *PHAS* loci accumulate 21-nt phasiRNAs with peak abundance at the premeiotic stage of anthers (83.2% to 99.6%), similar to those of maize and rice (Zhai et al. 2015; Fei et al. 2016). Although most Pooideae *PHAS* loci produce 21-nt phasiRNAs that peak in premeiotic anthers, we observed distinct groups of *PHAS* loci generating phasiRNAs with peak accumulation at the meiotic and postmeiotic stages in sampled species. For example, in oat, while the majority of 21-*PHAS* loci produced phasiRNAs during the premeiotic stage (9,007 out of 10,044; 89.7%), a substantial number of loci also produced phasiRNAs during the meiotic (660 out of 10,044; 6.6%) and postmeiotic (377 out of 10,044; 3.8%) stages (Figure 2A, Supplemental Table 4). Similar patterns were observed in anthers of barley, rye, and bread wheat (Supplemental Figure 3A). Meanwhile, in *Brachypodium* and durum wheat, we detected groups of 21-nt phasiRNAs primarily during the premeiotic and postmeiotic stages (Figure 2A). No significant differences were found in the median abundance of phasiRNAs peaking at the three developmental stages in all species. However, it’s noteworthy that in oats, we observed two 21-*PHAS* loci that accumulated exceptionally abundant 21-nt phasiRNAs (>5000 reads per million mapped, RPMM) in postmeiotic anthers, even though the median abundance was 394.2 RPMM. In most species, postmeiotic anthers displayed extremely abundant 21-nt phasiRNAs from a small group of 21-*PHAS* loci. No homology of these loci was observed among sampled species.

**Figure 2:**
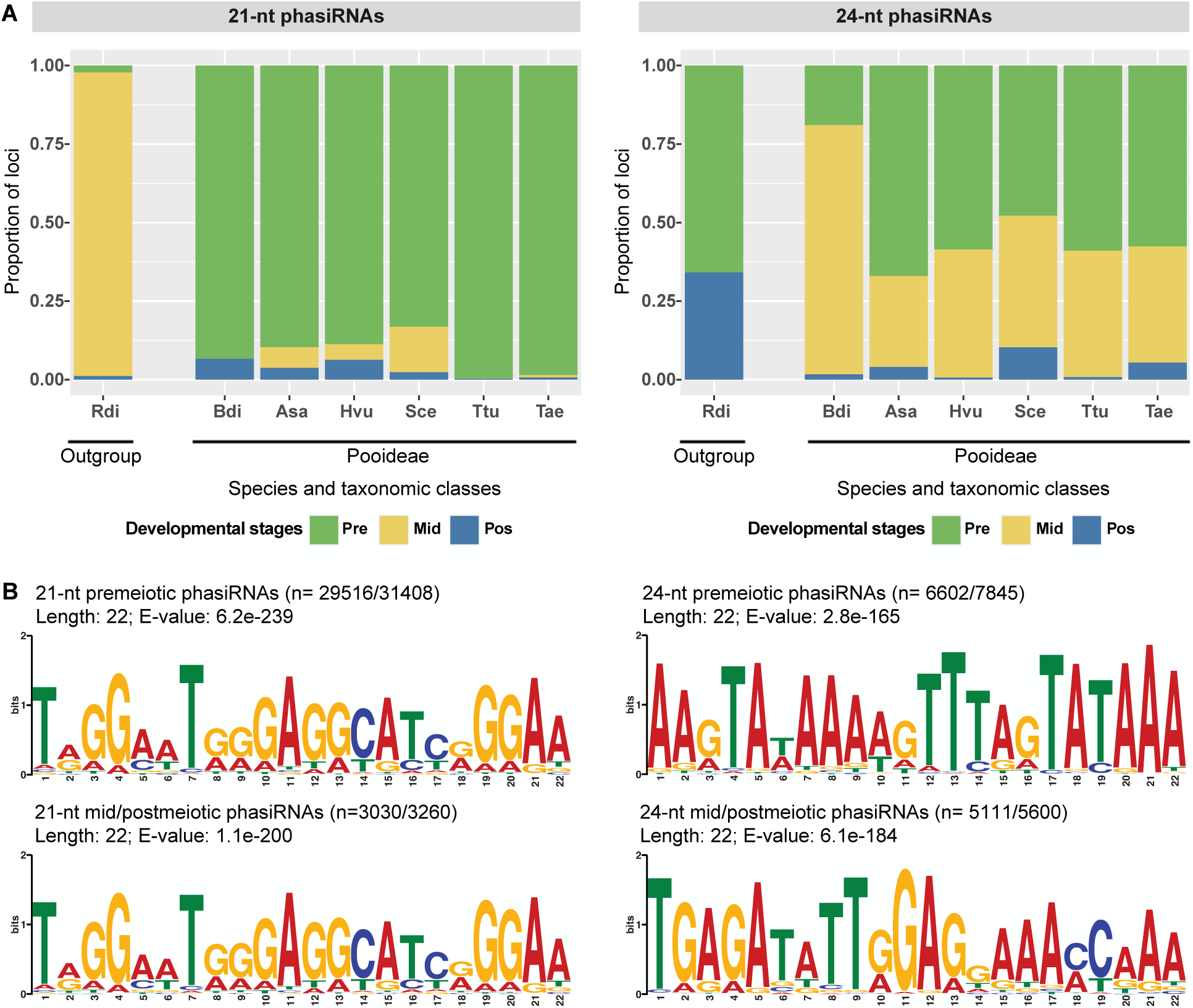
Reproductive phasiRNAs are tightly regulated throughout developmental stages. (A) A stacked bar chart illustrates the proportion of *PHAS* loci producing 21-nt (on the left) and 24-nt (on the right) phasiRNAs, accumulating during premeiotic, meiotic, and postmeiotic stages in seven species. The graph demonstrates a distinct accumulation profile of reproductive phasiRNAs in developing anthers of bamboo compared to Pooideae species. Notably, we observed that groups of 21-*PHAS* loci are producing phasiRNAs not only during the premeiotic stage of anther development in multiple species but also that groups of 24-*PHAS* loci express phasiRNAs during the premeiotic stage of anther development in all species. (B) Conserved motifs were identified in putative 21-*PHAS* (on the left) and 24-*PHAS* (on the right) precursor transcripts of premeiotic (upper) and meiotic/postmeiotic (lower) phasiRNAs. The miR2118 motif was enriched in all 21-*PHAS* precursor transcripts. However, the miR2275 motif was found exclusively in meiotic and postmeiotic 24-*PHAS* precursors, while a distinct enriched motif identified in premeiotic 24-*PHAS* precursors did not match any known miRNAs or sRNAs sequenced in our libraries. We use three-letter codes to represent the seven species as follows: bamboo (*R. distichophylla*; Rdi), oat (*A. sativa*; Asa), *Brachypodium* (*B. distachyon*; Bdi), barley (*H. vulgare*; Hvu), rye (*S. cereale*; Sce), durum wheat (*T. turgidum*; Ttu), and bread wheat (*T. aestivum*; Tae).

Premeiotic 21-nt phasiRNAs require a specialized miRNA trigger, miR2118, in rice and maize (Komiya et al. 2014). Therefore, we searched for conserved motifs in putative *PHAS* precursors to determine if the miR2118 motif is conserved among species and groups of premeiotic and meiotic/postmeiotic 21-*PHAS* loci. Notably, considering all sampled species, we found that most premeiotic (29,516 out of 31,408; 94.0%) and meiotic/postmeiotic (3,030 out of 3,260; 93.0%) 21-*PHAS* loci have a 22-nt conserved motif homologous to miR2118 (Figure 2B). Analysis of each species separately found the same results. Several loci expressed miR2118 in the genomes of all sampled species. We found miR2118 expression at corresponding developmental stages when 21-nt phasiRNAs accumulate in all seven species (Supplemental Table 3), suggesting a conserved role of miR2118 in triggering the cleavage of 21-*PHAS* precursors regardless of developmental stage or species.

### Temporal expression and biogenesis of 24-nt phasiRNAs in anthers diverged across Poaceae subfamilies

While the majority of *PHAS* loci generate 24-nt phasiRNAs primarily during the meiotic stage in maize and rice anthers, they are predominant at the premeiotic stage in barley and bread wheat (Bélanger et al. 2020; Zhan et al. 2024). Here, we investigated the evolution of the temporal accumulation patterns of 24-nt phasiRNAs in the Pooideae and Bambusoideae. In the Pooideae, we found distinct groups of 24-*PHAS* loci generating phasiRNAs that peak at each of the three developmental stages. In oat, barley, and the two wheat species, the premeiotic stage displayed the greatest number of 24-*PHAS* loci (oat: 2,284 out of 3,408, 67.0%; barley: 472 out of 806, 58.6%; durum wheat: 1,610 out of 2,731, 59.0%; bread wheat: 2,457 out of 4,264, 57.6%) followed by meiotic (oat: 987, 29.0%; barley: 329, 40.8%; durum wheat: 1,100, 40.3%; bread wheat: 1,577, 37.0%) and postmeiotic (oat: 137, 4.0%; barley: 5, 0.6%; durum wheat: 21, 0.8%; bread wheat: 230, 5.4%) stages (Figure 2A, Supplemental Figure 3B and Supplemental Table 5). However, in rye, about equal numbers of 24-*PHAS* loci are generating phasiRNAs at the premeiotic (829 out of 1,733, 47.8%) and meiotic (726, 41.9%) stages, followed by the postmeiotic stage (178, 10.3%). Additionally, the temporal accumulation pattern of 24-nt phasiRNAs in *Brachypodium* is similar to those in rice and maize (Zhan et al. 2024), with the meiotic stage as the main stage generating phasiRNAs from 24-*PHAS* loci (234 out of 295, 79.3%) (Figure 2A, Supplemental Figure 3B and Supplemental Table 5). The temporal accumulation of 24-nt phasiRNAs in anthers of bamboo differ from the six Pooideae species as well as maize and rice. The 24-*PHAS* loci do not accumulate phasiRNAs at the meiotic stage at all, while they are mainly observed at premeiotic (137 out of 208, 65.9%), followed by postmeiotic stage (71, 34.1%). (Figure 2A, Supplemental Figure 3B and Supplemental Table 5). Together, these results indicate divergent temporal abundance patterns of 24-nt phasiRNAs across Poaceae subfamilies, potentially indicating different roles in anther development.

The biogenesis of meiotic 24-nt phasiRNAs in maize and rice requires a specialized miRNA trigger, miR2275 (Zhai et al. 2015; Fei et al. 2016). We found that most meiotic/postmeiotic (5,111 out of 5,600; 91.2%) 24-*PHAS* loci possess a 22-nt conserved motif homologous to miR2275 and the motif is conserved in Pooideae species (Figure 2B and Supplemental Figure 4). While meiotic 24-nt phasiRNAs are absent in bamboo, it is notable that *PHAS* precursors of postmeiotic 24-nt phasiRNAs exhibit the exact same conserved motif homologous to miR2275 (Supplemental Figure 4) suggesting conserved biogenesis mechanisms despite being generated at a different developmental stage. Several loci in the genomes of all sampled species produce mature miR2275 and we found miR2275 expression at corresponding developmental stages when meiotic and postmeiotic 24-nt phasiRNAs accumulate in sampled species (Supplemental Table 3), suggesting a conserved role of miR2275 in triggering the cleavage of meiotic and postmeiotic 24-*PHAS* precursors.

In contrast to the conserved meiotic 24-nt phasiRNAs precursor in all studied Poaceae species, the predicted motifs for premeiotic 24-*PHAS* loci differ among clades and species. All Triticeae species (barley, rye, durum wheat and bread wheat) and oat share the same 22-nt motif among most of the premeiotic 24-*PHAS* loci, while *Brachypodium* and bamboo have their own unique motifs (Supplemental Figure 4). Although the same 22-nt motif was conserved among most premeiotic 24-*PHAS* loci in oat (1,900 out of 2,284, 83.2%) and the four Triticeae (barley: 388 out of 472, 82.2%; rye: 501 out of 829, 60.4%; durum wheat: 1,365 out of 1,610, 84.8%; bread wheat: 1,323 out of 2,457, 53.8%), none of the known miRNAs match this motif, and we could not identify any sRNA among the 63 libraries sequenced in this work that matched it either (Figure 2B; Supplemental Figure 4). Similarly, *Brachypodium* (43 out of 56; 76.8%) and bamboo (53 out of 137; 38.0%) premeiotic 24-*PHAS* precursors exhibit alternative and unique motifs that do not correspond to any known miRNA or other sRNA sequences identified in this study (Supplemental Figure 4). None of the motifs described above match the conserved motifs identified in maize and teosinte (Zhan et al. 2024). This suggests that the biogenesis of premeiotic 24-nt phasiRNAs evolved independently across Pooideae subfamilies.

### Positional and compositional nucleotide signatures are observed among phasiRNAs

We analyzed the frequency of nucleotides to capture positional and compositional signals along 21-nt and 24-nt phasiRNAs. PhasiRNAs generated at premeiotic and meiotic/ postmeiotic stages of anthers were analyzed separately. To calculate the nucleotide frequency at each position within the phasiRNAs, we merged the data from all species. We retrieved all sRNAs from all *PHAS* loci, producing from 8.9 M to 85.4 M data points, and calculated the nucleotide frequency at each position (Supplemental Figure 5). We found clear compositional biases at specific positions along 21-nt and 24-nt phasiRNAs, while positions with no signal showed a neutral 0.25 frequency. Among 21-nt phasiRNAs, we found that positions 1, 3, 19 and 21 were overrepresented by C/G nucleotides while positions 2, 8, 14 and 20 were overrepresented by A/U. For 24-nt phasiRNAs, we found that positions 1, 2, 23 and 24 were overrepresented by A/G, A, U and U/C nucleotides, respectively. The compositional signals that best differentiated the 21-nt and 24-nt phasiRNAs were at positions 3, 8 and 14 at which specific nucleotides were overrepresented among 21-nt phasiRNAs, whereas the frequencies were neutral at these positions in 24-nt phasiRNAs. These biased nucleotides and positions may influence the loading of phasiRNAs in AGO effectors.

### Annotation and expression analyses of key genes involved in phasiRNA biogenesis and function

We annotated the key genes involved in phasiRNA pathways, including *AGO*, *DCL* and *RDR*, and performed their phylogenetic and co-expression analyses (Supplemental Figure 6; Supplemental Table 6 to 8). Specific *AGO* genes, namely, *AGO4b*, *AGO5a/e*, and *AGO7,* are not expressed in anthers of most sampled species (Figure 3). Most of the expressed *AGO* genes were differentially abundant over the course of developing anthers (Figure 3 and Supplemental Table 8). The accumulation of stage-specific reproductive phasiRNAs corresponded with specific and distinct *AGO* genes. Across species, *AGO1b/d*, *AGO5c/d*, *AGO4a/c*, and AGO10 were coexpressed with the premeiotic anthers, as the expression of these *AGOs* was gradually downregulated from premeiotic to postmeiotic anther stages (Figure 3 and Supplemental Table 8). These *AGO* genes were coexpressed with two groups of phasiRNAs, namely the premeiotic 21-nt and 24-nt phasiRNAs. Accumulation of groups of meiotic/postmeiotic 21-nt and 24-nt phasiRNAs was concurrent with *AGO2b*, *AGO5b*, *AGO6*, and *AGO18*. Notably, orthologous genes annotated across the seven species exhibited the same expression profiles, indicating a strong association between the genes mentioned above and anther developmental stages. Moreover, multiple copies of the same gene showed identical expression patterns in the polyploid wheat and oat genomes. For example, the bread wheat genome encodes a total of 10 AGO5c/d proteins for which nine genes were expressed in anthers and all the corresponding genes were coexpressed in the premeiotic stage. Clades of functionally characterized rice *AGO* genes in the phasiRNA pathways are annotated on Figure 3. Candidate *AGO* genes with a potential function in the phasiRNA pathways were annotated in the tree.

**Figure 3:**
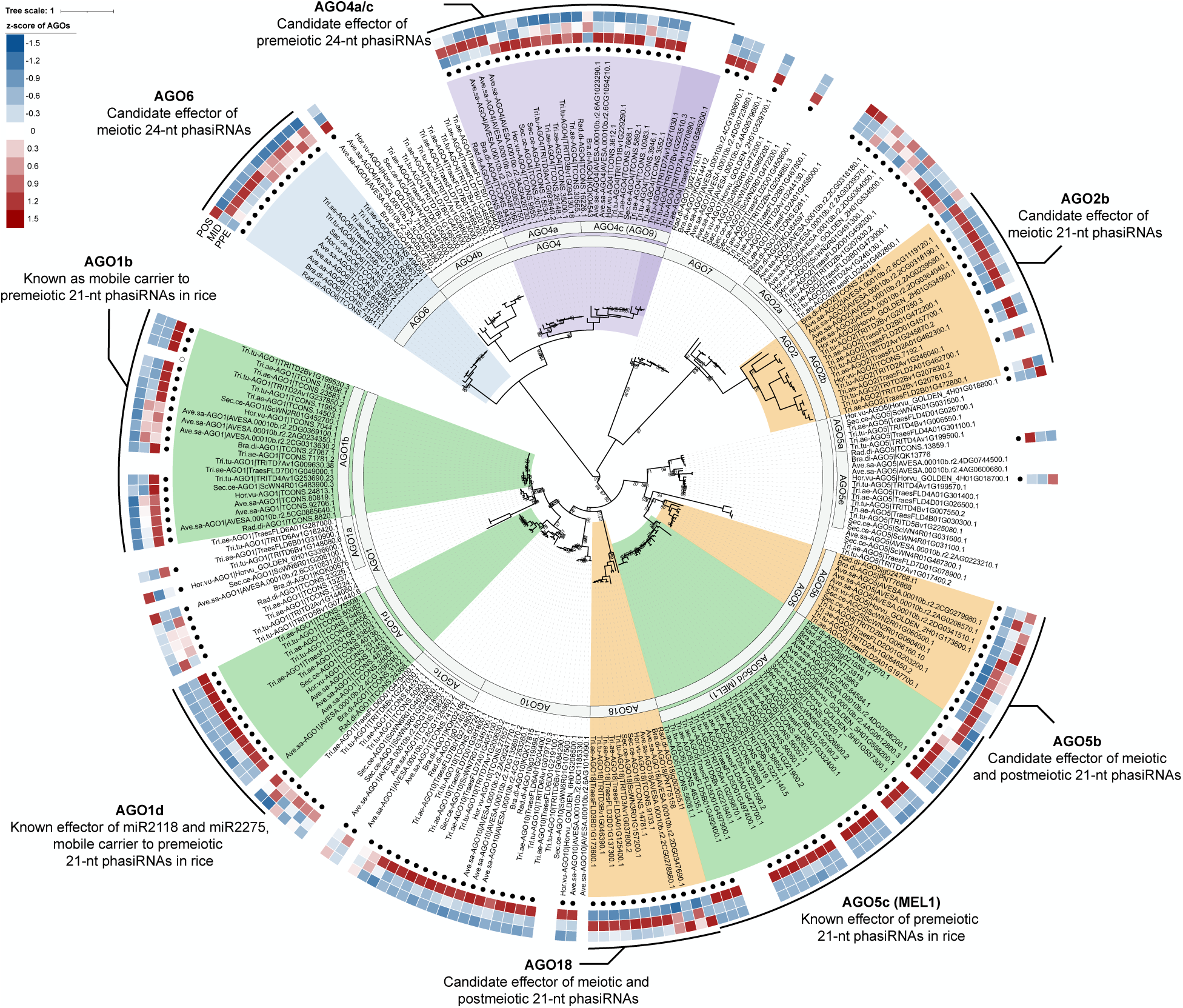
The phylogenetic tree and expression profiles of genes encoding AGO proteins in the sampled species. The phylogenetic tree was derived from the maximum-likelihood analysis based on 39 species. Clades representing AGO proteins are clearly indicated. Genes expressing AGO proteins in anthers are marked with a black dot and the heatmap denoting the calculated z-score relative expression change between premeiotic (inside), meiotic (middle) and postmeiotic (outside) stages. Protein clades with known or predicted functions in phasiRNA biogenesis and function are also highlighted.

Almost all *DCL* and *RDR* genes were expressed in anthers of all species, however, a lower number of *DCL* and *RDR* genes were differentially expressed over the course of developing anthers, and the shared expression profile between species is less clear than for the *AGOs*. For instance, *DCL4* was not significantly differentially expressed but had an expression pattern that decreases gradually from premeiotic to postmeiotic anthers, consistent with the accumulation pattern of 21-nt phasiRNAs. *DCL5* genes were extremely abundant and were differentially expressed in almost all species. The abundance of *DCL5* transcripts peaked at the premeiotic (*Brachypodium* and durum wheat) or meiotic (oat, bamboo, barley, and bread wheat) stages in most species which coincides with the two groups of 24-nt phasiRNAs (Supplemental Figure 7A and 8). However, the peak abundance of *DCL5* at the postmeiotic stage in rye did not correlate with peaks of 24-nt phasiRNA accumulation. Duplicated in oat and the Triticeae (barley, rye and wheat species), *RDR1* paralogs were differentially expressed in all species, with the duplicated *RDR1* copies exhibiting contrasting expression patterns with peaks of abundance in premeiotic or postmeiotic stages (Supplemental Figure 7B and 8). *RDR6* genes were differentially expressed in the Triticeae (barley, rye, durum wheat and bread wheat) but not in *Brachypodium*, oat or bamboo (Supplemental Figure 7B and 8). The gradual downregulation of *RDR6* from premeiotic to postmeiotic anthers correlates with the peak of abundance of 21-nt and 24-nt phasiRNAs observed across species.

### Degradome sequencing revealed little overlap among miRNA and phasiRNA targets in anthers across species

To assess the post-transcriptional activity of miRNAs and phasiRNAs in developing anthers, we performed nanoPARE analyses to (i) validate the cleavage sites of miRNA target genes and (ii) to identify targets of phasiRNAs. We used PAREsnip2 (Thody et al. 2018) to predict and validate target cleavage sites (category ≤ 1) for the distinct miRNAs annotated among species. We validated a total of 21 to 129 genes to be targets of miRNA families annotated in anthers of the seven species (Supplemental Table 9 and 10). Supplemental Table 9 shows the proportion of targeted genes having an open reading frame (ORF) or not. Targeted genes with no ORF are essentially the targets of miR2118 and miR2275, which are the two miRNA families known to trigger the biogenesis of 21-nt and meiotic 24-nt phasiRNAs. These target transcripts overlap with the few *PHAS* precursors retrieved in the transcript assemblies mentioned in the previous section. To evaluate the extent of overlap between protein-coding transcripts targeted by miRNAs across species, we retrieved the orthogroups (i.e. groups of genes descended from a single gene in the last common ancestor of a group of species) containing the targets. Figure 4A shows the number of orthogroups representing the targeted protein-coding genes across species and their overlap among species. Notably, we found three orthogroups present in all seven species. These are homologous genes corresponding to known targets of four miRNAs, namely, miR166-*HOMEOBOX-LEUCINE ZIPPER PROTEIN* (*HD-ZIP*), miR167-*AUXIN RESPONSE FACTORs* (*ARF*), and miR396-*GROWTH REGULATING FACTORS* (*GRF*). Taking barley as an example, we validated cleavage sites for multiple paralogs for *ARF* (orthologous to *OsARF8*, *OsARF10*, *OsARF12*, *OsARF17*, *OsARF18*, *OsARF22* and *OsARF25*), *GRF* (orthologous to two *OsGRF3*, *OsGRF4*, *OsGRF6*, *OsGRF10* and *OsGRF12*), and *HD-ZIP* (orthologous to *OsHOX32* and *OsHOX33*) genes. Target cleavage tags, as well as the corresponding miRNAs, were found to overlap between replicates, but not between all developmental stages, suggesting that most replicates have good consistency in the degradome data and that some of these contribute to explaining differences between stages (Supplemental Table 10). Notably, we found that most miRNA targets were found in a single species (Figure 4A), suggesting species-specific miRNA activities over the course of anther development.

**Figure 4:**
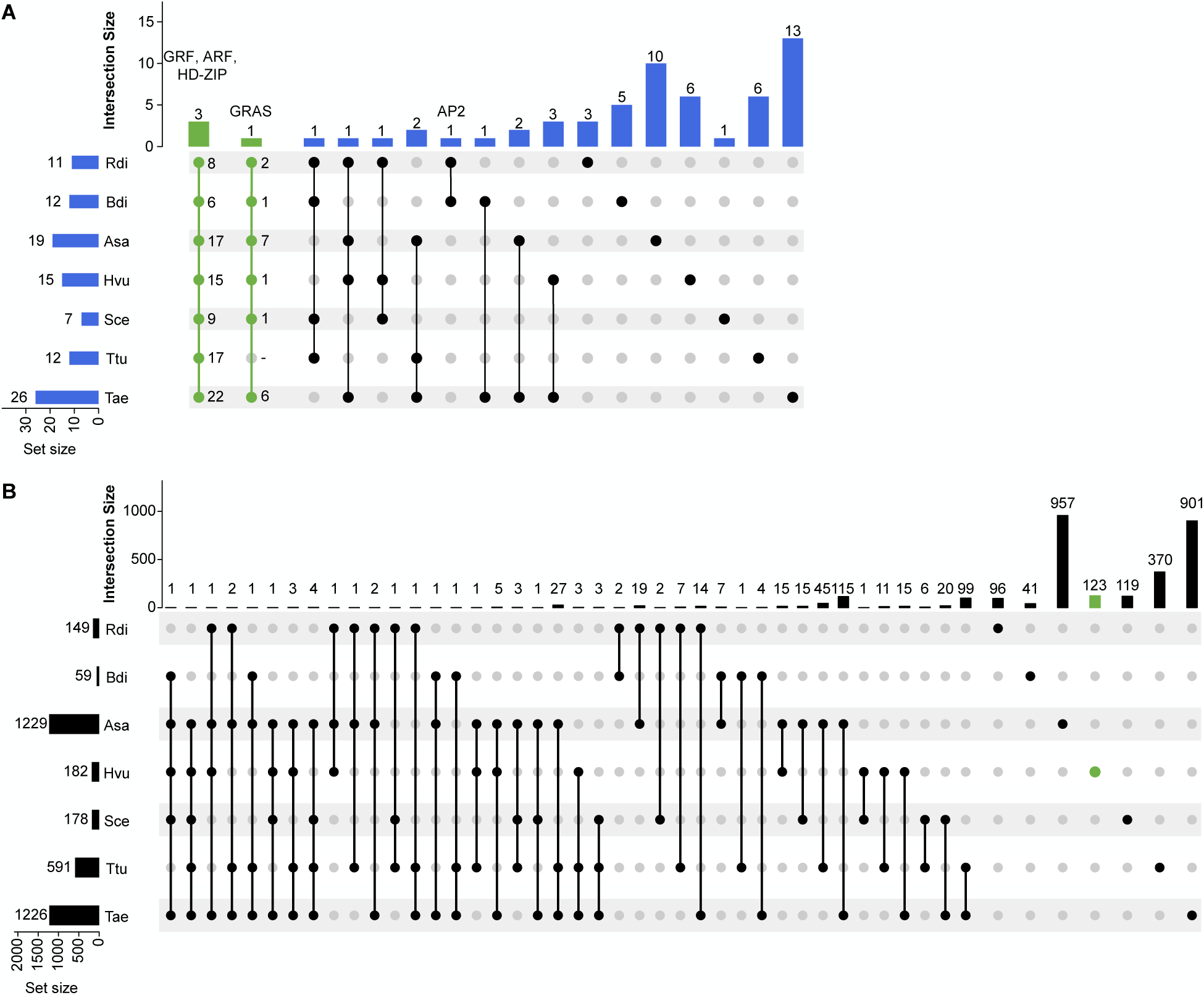
Degradome sequencing revealed little overlap among miRNA and 21-nt phasiRNA targets in anthers across species. Upset plots illustrate orthogroups comprising protein-coding genes targeted by miRNA (A) and 21-nt phasiRNA (B) shared among different species. The bottom-left plot represents the size of each set with a horizontal histogram, the bottom-right displays the intersection matrix, and the upper-right indicates the size of each combination with a vertical histogram. Intersection data selected for discussion in the results section are highlighted in green. We use three-letter codes to represent the seven species as follows: bamboo (*R. distichophylla*; Rdi), oat (*A. sativa*; Asa), *Brachypodium* (*B. distachyon*; Bdi), barley (*H. vulgare*; Hvu), rye (*S. cereale*; Sce), durum wheat (*T. turgidum*; Ttu), and bread wheat (*T. aestivum*; Tae).

Our nanoPARE analyses of 21-nt phasiRNAs validated a total of 70 to 1,983 genes as targets of this phasiRNA class in anthers of the seven species (Supplemental Table 9 and 11). Supplemental Table 9 shows that most targeted genes encode proteins. Thus, we retrieved the orthogroups containing the 21-nt phasiRNA targets to evaluate the extent of overlap between species and found almost no overlap (Figure 4B). Only 14 orthogroups overlapped in four species or more from a total of 3,070 orthogroups represented by 4,496 targeted protein encoding genes, suggesting species-specific 21-nt phasiRNA activities over the course of anther development rather than a conserved function across species (Figure 4B and Supplemental Table 9). The detailed results are shown in Supplemental Table 11. While minimal overlap was observed across species, we concentrated on observations in barley: a total of 185, 33, and 77 protein-coding targets, respectively, were validated at the premeiotic, meiotic, and postmeiotic stages. All targets validated in both meiotic and postmeiotic stages were also detected at the premeiotic stage, indicating that the cleavage of 21-nt phasiRNAs was initiated in premeiotic anthers coinciding with the burst of phasiRNAs. Among the targeted genes, we identified genes encoding proteins with molecular or biological functions such as structural constituents of chromatin (orthologous to Histone H4), chromatin remodeling (orthologs of *AtDRD1*, *AtSWI3B*, *AtRHL1*), meiosis (orthologs of *AtHOP2* and *OsEME1*), transcription factors (including orthologs of *AtbHLH105*, *OsbHLH3*, *OsGRAS-38*, *AtMGP*, and *OsSCL7*), regulation of transcription (ortholog of *AtDRD1*), post-transcriptional gene silencing (orthologs of *AtXRN3* and *OsDRB5*), and regulation of translation (orthologs of *AtENODL14*, *OsRACK1A*, and *OsRPL9*). We identified the cleavage of an orthologous gene to the rice *MULTIPLE SPOROCYTE 1* (*OsMSP1*) which is known to play important roles in restricting the number of cells entering into male and female sporogenesis. In barley, cleavage events were detected in premeiotic anthers for these genes. The downregulation of target genes in later developmental stages, especially postmeiotic stages, suggests that 21-nt phasiRNAs play a role in coordinating the developmental transition (Supplemental Table 11).

In contrast to the large number of genes validated as targets of 21-nt phasiRNAs, only 4 to 120 genes were validated for cleavage by 24-nt phasiRNAs, with no overlap among species (Supplemental Table 9 and 12). These findings suggest that either the majority of reproductive 24-nt phasiRNAs function through mechanisms other than mediating mRNA cleavage, or they target only a small subset of genes that have rapidly diverged across Pooideae species.

## DISCUSSION

In this work, we staged and selected anthers at premeiotic, meiotic and postmeiotic stages for deep transcriptomic profiling to investigate the evolution, conservation, regulation, and function of 21-nt and 24-nt phasiRNAs in one bamboo and six Pooideae (*Brachypodium*, oat, rye, barley, durum and bread wheat) species. The current work and a previous finding (Zhan et al. 2024) together suggest that premeiotic 24-nt phasiRNAs are quite broadly present in the grass family as their presences had been confirmed across species in Panicoideae, Oryzoideae, Bambusoideae, and Pooideae subfamilies. We found that premeiotic and meiotic 24-nt phasiRNAs are similar in their nucleotide composition, with conserved nucleotides at 5’-end and 3’-end positions. The precursor of meiotic 24-nt phasiRNAs in all analyzed grass species contains the miR22775 motif; the premeiotic 24-*PHAS* precursors trigger sites are conserved in Triticeae and oat, but differ across grass subfamilies (Zhan et al. 2024). None of the known miRNAs match these motifs and we could not find any matching sRNAs among sequenced libraries, suggesting that premeiotic 24-nt phasiRNAs are not initiated via a miRNA-directed, AGO-catalyzed cleavage of single-stranded *PHAS* precursor, but by an as-yet unknown mechanism.

Notably, our results show that the accumulation patterns of both 21-nt and 24-nt phasiRNAs differ between bamboo and any studied grass species, including the six Pooideae species, maize, teosinte and rice (Zhai et al. 2015; Fei et al. 2016; Zhan et al. 2024). In bamboo, almost all 21-*PHAS* loci accumulate at the meiotic stage of anthers rather than the premeiotic stage as observed in the Pooideae species (Figure 2A). An analysis of putative 21-*PHAS* precursors revealed that most premeiotic, meiotic and postmeiotic precursors have a 22-nt conserved motif homologous to miR2118, suggesting that 21-nt phasiRNA biogenesis is triggered by the same *MIR* family and thus their temporal accumulation may be related to a stage-specific activation of 21-*PHAS* precursor transcripts (Figure 2A). Regarding 24-nt phasiRNAs, no *PHAS* loci produce meiotic 24-nt phasiRNAs in bamboo while there are hundreds to thousands of meiotic 24-*PHAS* loci in Pooideae species, maize, teosinte, and rice. Moreover, we found 34.1% postmeiotic 24-*PHAS* loci in bamboo, of which 86.0% have the miR2275 trigger motif, while fewer (0.6% to 10.3% 24-*PHAS* loci) were annotated at the postmeiotic stage in the Pooideae species. Considering that miR2275-triggered meiotic 24-nt phasiRNAs are widely conserved among angiosperms (Xia et al. 2019), these observations indicate significant variations in the temporal accumulation of reproductive 24-nt phasiRNAs between Bambusoideae and other species during anther development in meiotic and postmeiotic stages.

In rice and maize, multiple AGO proteins are involved. AGO1d catalyzes the cleavage of 21-*PHAS* precursors (Shi et al. 2022). AGO1b and AGO1d are mobile carriers of 21-nt phasiRNAs from epidermal cells to sporocytes (Tamotsu et al. 2023). AGO1d, AGO5b (MALE-ASSOCIATED ARGONAUTE-1, ZmMAGO1 in maize) and AGO5c (MEIOSIS ARRESTED AT LEPTOTENE 1, OsMEL1, ZmMAGO2) are effectors of 21-nt phasiRNAs in developing meiotic cells (Komiya et al. 2014; Shi et al. 2022). In this study, we observed that the expression of *AGO1b*, *AGO1d* and *AGO5c* genes coincide with peak accumulation of most 21-nt phasiRNAs in premeiotic anthers of the Pooideae suggesting that these AGOs have a conserved function with maize and rice in the phasiRNA biogenesis and function. In bamboo, there are no premeiotic 21-nt phasiRNAs as these accumulate in the meiotic stage of anthers. Interestingly, *AGO1d* and *AGO5c* are expressed in meiotic anthers which, in bamboo, corresponds to the peak abundance for most 21-nt phasiRNAs (Figure 2A and 3). We described 21-*PHAS* loci producing phasiRNAs in meiotic and postmeiotic anther in Pooideae and we found that all species express *AGO2b*, *AGO5b* and *AGO18* that peak at the same developmental stage. *AGO5b* seems to be a good candidate effector gene for meiotic and postmeiotic 21-nt phasiRNAs as *AGO5b* is orthologous to the maize *MAGO1* gene and proposed to be functionally redundant with *AGO5c* to catalyze the function of pre-meiotic 21-nt phasiRNA in maize (Lee et al. 2021).

We observed that the expression of *AGO4a/c* and *AGO6* genes coincide with a burst of premeiotic and meiotic 24-nt phasiRNAs, respectively. The expression of *AGO4a/c* genes synchronizes with the premeiotic stage of anthers of most species. The same peak of expression was observed for *AGO6* genes in meiotic anthers of all Pooideae species. In the absence of meiotic 24-nt phasiRNAs in bamboo, *AGO6* abundance peaks in postmeiotic anthers when 24-nt phasiRNAs accumulate. The conserved expression of *AGO4a/c* and *AGO6* with premeiotic and meiotic anthers across the species, together with the enriched nucleotides at the 5’-end of 24-nt phasiRNAs, strengthens the hypothesis that they may be binding partners to 24-nt phasiRNAs. In summary, most orthologous genes involved in phasiRNA biogenesis and function exhibited the same expression profiles in species with similar phasiRNA accumulation patterns, indicating a strong association between the genes mentioned above and anther developmental stages.

Our degradome analyses reveal conservation of targets among three miRNA families across species. However, the majority of miRNA targets are species-specific, indicating unique miRNA activities during anther development. Moreover, although the 21-nt phasiRNAs mediate cleavage of a notable number of target mRNAs, we detected limited orthologous targets, suggesting that the 21-nt phasiRNA target diverged rapidly. On the other hand, the numbers of 24-nt phasiRNA targets are very small, with no orthologous targets, suggesting that the majority of 24-nt phasiRNAs do not mediate mRNA cleavage. This may be attributable to the function of meiotic 24-nt phasiRNAs in CHH DNA methylation (Zhang et al. 2021), or the 24-nt phasiRNA targets being cell type/layer specific so that they are not detected by nanoPARE analysis performed on intact anthers.

In summary, this work shows that premeiotic 24-nt phasiRNAs are present across Bambusoideae and Pooideae species but that the temporal accumulation of other classes of 21-nt and 24-nt phasiRNA differs between bamboo and Pooideae species. The targets of the three classes of phasiRNAs are likely rapidly evolving or not yet detected, perhaps requiring single-cell methodologies.

## MATERIAL AND METHODS

### Plant growth conditions and tissue harvesting

We grew *Brachypodium* (*Brachypodium distachyon* ‘Bd21-3’), oat (*Avena sativa*, ‘Fraser’), barley (*Hordeum vulgare ssp. vulgare*, ‘Golden Promise’), rye (*Secale cereale*, ‘Horton’), durum wheat (*Triticum turgidum subsp. durum*, ‘Kronos’), and bread wheat (*Triticum aestivum*, ‘Fielder’) plants in a greenhouse under the following conditions: 20°C/18°C day/night temperature, 16 hours of light and 8 hours of darkness, and 50% relative humidity. Herbaceous bamboo (*Raddia distichophylla*) plants were grown in a chamber under shade cloth, with conditions set at 26°C/18°C day/night temperature, 12 hours of light and 12 hours of darkness, and 75% relative humidity.

We used a stereomicroscope (Mantis Elite-Cam HD, Vision Engineering) to dissect anthers. The length of the anthers collected for staging varied between species. For *Brachypodium*, we collected anthers ranging from 0.1 mm to 0.8 mm with 0.1 mm increments. For the other six species, we collected anthers at 13 different lengths: 0.2, 0.4, 0.6, 0.8, 1.0, 1.2, 1.4, 1.6, 1.8, 2.0, 2.5, 3.0, and 3.5 mm. Additional samples were harvested for oat and rye to collect anthers measuring 4.0 and 5.0 mm in size for oat and 4.0, 5.0, 7.5, 10.0, and 12.5 mm for rye. For each stage, we harvested at least 10 anthers, which were promptly fixed with aldehyde for histological experiments. For RNA experiments, we collected anthers at premeiotic, meiotic, and postmeiotic stages. For each stage, we harvested 15, 30, and 50 anthers per sample, respectively. The corresponding lengths of the anthers harvested at each stage are indicated in Figure 1D. We performed three replicates for anther harvesting, and after dissection, the samples were immediately frozen in liquid nitrogen and stored at −80°C until RNA isolation.

### Anther staging

We prepared samples as described in Bélanger et al. (Bélanger et al. 2020, 2022) with modifications. Briefly, freshly harvested anthers were fixed overnight in a solution containing 2% [v/v] paraformaldehyde, 2% [v/v] glutaraldehyde, and 0.1% [v/v] Tween20 in 0.1 M PIPES buffer at pH 7.4. Subsequently, the anthers were dehydrated through a standard acetone series (30%, 50%, 70%, 80%, 90%, and 100% [v/v] cold acetone). Following dehydration, the samples were infiltrated and embedded in the Quetol 651 - NSA Kit (no. 14640, Electron Microscopy Sciences) and polymerized at 60°C. The embedded tissues were then sectioned at 500 nm using the Leica Ultracut UCT (Leica Microsystems Inc., Wetzlar, Germany). Sections were stained using the Epoxy Tissue Stain solution (no. 14950, Electron Microscopy Sciences) and mounted on a slide for image capture. Anther sections were observed using a Leica DM 750 microscope, and images were captured using a Leica ICC50 HD camera along with Leica Acquire v2.0 software (Leica Microsystems Inc., Wetzlar, Germany). The images were subsequently analyzed using ImageJ (Schindelin et al. 2012).

### RNA isolation, library construction, and sequencing

Total RNA was isolated using the TRI Reagent (Sigma-Aldrich, St. Louis, USA) according to the manufacturer’s instructions. The quality of RNA was assessed using an Agilent RNA 6000 Nano Kit on the Bioanalyzer 2100 (Agilent Technologies, Santa Clara, USA). Only samples with an RNA integrity number (RIN) greater than 7.0 were selected for library construction. We used the RealSeq-AC miRNA Library Kit for Illumina sequencing (Somagenics, Santa Cruz, USA) to construct the sRNA libraries. A total of 150 ng of RNA was used as input and 16 PCR amplification cycles were performed. The sRNA libraries were size-selected for an end product of 150 nt using the SPRIselect Reagent (Beckman Coulter Life Sciences, Indianapolis, USA) magnetic beads. We prepared RNA-Seq (Smart-seq2; Illumina, San Diego, USA) and nanoPARE libraries from 5 ng total RNA according to the nanoPARE library preparation protocol using the nanoPARE library preparation protocol (Schon et al. 2018) following modifications described by Pokhrel *et al*. (Pokhrel et al. 2021). Single-end reads were generated with 76 cycles for sRNA, nanoPARE and mRNA libraries using an Illumina NextSeq 550 instrument (Illumina, San Diego, USA) at the University of Delaware DNA Sequencing and Genotyping Center.

### Preprocessing and mapping of sRNA-Seq data

We utilized cutadapt v4.1 (Martin 2011) to preprocess sRNA-Seq reads. The preprocessing steps included the removal of the first 5’ nucleotide (-u 1), the 3’ adapter (-a TGGAATTCTCGGGTGCCAAGG), and the reads outside the length range of 19-nt to 25-nt (--minimum-length 19 to --maximum-length 25). We employed ShortStack v4.0.2 (Johnson et al. 2016) to map preprocessed sRNA reads to reference genome assemblies (Supplemental Table 1). The parameters used for mapping were as follows: --threads 20 --mmap u –mincov 1 --align_only --dicermin 20 --dicermax 24 --pad 75. Following the read mapping, we conducted an analysis of the mapped reads to annotate phasiRNAs and miRNAs.

### Annotation of miRNAs

We used ShortStack v4.0.2 (Johnson et al. 2016) to analyze alignment files and identify hairpin-derived sRNA loci. Predicted miRNA loci meeting the criteria for plant miRNA annotations (Axtell and Meyers 2018) were selected for downstream analyses. The criteria for true positive miRNAs were as follows: (i) identification of a precursor, (ii) presence of the miRNA* read, and (iii) mapping of over 75% of reads to the miRNA/miRNA* duplex. To annotate known miRNA families, we aligned representative reads of miRNA loci (both miRNA and miRNA*) to monocot-derived miRNAs listed in miRBase release 22.1 (Kozomara and Griffiths-Jones 2014; Kozomara et al. 2018) using ncbi-blastn v2.11.0+ (Camacho et al. 2009) with the following parameters: -strand both -task blastn-short -perc_identity 85 -word_size 7 -evalue 0.01 -num_alignments 1 -no_greedy -ungapped. We filtered homology hits and classified sRNA reads as known miRNAs based on the following criteria: (i) no more than four mismatches and (ii) no more than 2-nt extension or reduction at the 5’ or 3’ end. It has been reported that miRNAs triggering 21-nt and 24-nt phasiRNAs, namely miR2118 and miR2275, are present at multiple loci in grass genomes, including polycistronic loci (Lan et al. 2022). These characteristics pose challenges in accurately mapping the mature miRNA and miRNA* to their true expression origins. The current mapping and miRNA annotation algorithms do not adequately address the resolution of the expressed miR2118 and miR2275 origins. Therefore, we did not consider the precision criteria for miRNA/miRNA* production and the hairpin structure as described by Axtell & Meyers (Axtell and Meyers 2018). Instead, we conducted a homology search using the predicted MIR2118 and MIR2275 precursor sequences to identify all potential loci expressing miR2118 and miR2275. In absence of a predicted miRNA hairpin structure, these loci were annotated as MIR2118 and MIR2275 loci if associated sRNAs map to these loci. miRNAs that did not show homology to known families were considered as previously unannotated miRNAs when the mature miRNA was 20-22 nt long. We used UpSetR v1.4.0 (Conway et al. 2017) to visualize overlap in miRNA families expressed in anther across species. We summarized known and previously unannotated miRNAs by family and renamed them based on genomic positions when multiple miRNA loci per family were present in the genomes. We compiled information on the coordinates, annotation and abundance of all miRNA loci (Supplemental Table 3).

### Annotations and analyses of phasiRNAs

We utilized ShortStack v3.8.5 (Johnson et al. 2016) to annotate loci that produce phasiRNAs. To identify genuine phasiRNA-producing loci, we considered sRNA clusters with a ‘Phase Score’ of 40 or higher and an abundance of 2.0 reads per million (RPM) or more. These loci are referred to as *PHAS* loci. The bioinfokit python toolkit (https://pypi.org/project/bioinfokit/) was employed to normalize the read count-matrix generated by ShortStack into Reads per Million Mapped (RPM). RPM-normalized reads were used to determine the peak of phasiRNA accumulation during the progression of meiosis and to visualize the relative abundance of 21-nt and 24-nt phasiRNAs using the R pheatmap v1.0.12 package (https://rdrr.io/cran/pheatmap/). *PHAS* loci were categorized based on the length of phasiRNAs produced (21-nt or 24-nt) and their peak accumulation during the premeiotic, meiotic, or postmeiotic stages. For each of the seven studied species, we compiled information on the coordinates, total abundance, peak accumulation (premeiotic, meiotic, or postmeiotic), and category (21-nt or 24-nt) of all annotated phasiRNA loci (Supplemental Table 5). We used the R package ggplot2 (Wickham 2016) to generate a stacked bar graph illustrating the proportion of PHAS loci accumulating at the three developmental stages across species.

We utilized the result report from ShortStack to assess the coordinates of *PHAS* loci and extract the putative precursors using bedtools slop with a -b value of 250 (Quinlan and Hall 2010). The putative precursors were then analyzed with MEME v5.4.1 (Bailey et al. 2015) to search and visualize conserved nucleotide motifs of premeiotic and meiotic/postmeiotic within putative 21-*PHAS* and 24-*PHAS* transcripts. We searched for motifs from *PHAS* precursors found in each species individually and from precursors merged from all species. Then, we analyzed the nucleic distribution at each position of premeiotic and meiotic/postmeiotic phasiRNAs expressed from 21-*PHAS* and 24-*PHAS* loci. This experiment was conducted separately for each species and for all species combined. In the first case, we simply collected the major sRNA read reported in the result file from ShortStack. In the second case, we used the BEDOPS v2.4.41 (Neph et al. 2012) functions, bam2bed with bedextract, to extract all sRNAs expressed within the coordinates of *PHAS* loci. For both cases, we visualized the data using the ggline function from ggpubr v.0.2.4.999 (Kassambara 2020).

### Analysis of RNA-Seq and nanoPARE data

We used cutadapt v4.1 (Martin 2011) to remove the adapters and trim low-quality nucleotides with a Phred quality score greater than 20 (-q 20). RNA-Seq and nanoPARE libraries were combined to assemble RNA transcripts, while nanoPARE libraries were analyzed separately to validate sRNA-directed cleavage sites.

To reconstruct transcripts expressed in anthers of each species, we mapped reads from RNA-Seq and nanoPARE libraries to the reference genomes using HISAT2 v2.2.1 (Kim et al. 2015) with default parameters. Genome-mapped reads were subjected to de novo reference-guided transcript assembly using two assemblers: StringTie v2.2.1 (-c 1.5, -f 0.2, -s 20, and -m 150) (Pertea et al. 2015, 2016) and Scallop v0.10.5 (--min_transcript_length_increase 35) (Shao and Kingsford 2017). Consensus transcripts were reconstructed using StringTie-merge with the following parameters: -F 1, -f 0.1, -m 200, and -g 1000 (Pertea et al. 2015, 2016). We quantified reads mapping to consensus transcript assemblies using Stringtie with the -e option, and we used the prepDE.py helper script to generate a gene-level read-count matrix. To denoise the matrix, we applied noisyR (Moutsopoulos et al. 2021) with the following parameters: similarity.measure = “correlation_spearman”, similarity.threshold = 0.20, method.chosen = “Boxplot-IQR”. The denoised matrix was filtered to remove genes with low abundance using edgeR (Robinson et al. 2010), resulting in a set of gene loci expressing transcripts at ≥ 3 cpm (counts per million reads). Gene loci expressing low-abundance transcripts were removed from the transcript assemblies. Differentially expressed genes (DEG) were identified using a threshold of absolute fold change of 1.5 premeiotic vs meiotic or meiotic vs postmeiotic or premeiotic vs postmeiotic, and adjusted P-value < 0.05, using DESeq2 (Love et al. 2014). The differentially expressed genes were analyzed using KOHONEN (Wehrens and Buydens 2007) for gene clustering. Supplemental Figure 8 summarizes the differentially expressed *AGO*, *DCL*, and *RDR* genes within coexpressed gene clusters. We used AGAT (https://github.com/NBISweden/AGAT) to extract expressed transcript sequences. Assembled GTF and transcript files were used to annotate known and novel protein sequences with TranSuite (Entizne et al. 2020). We used AGAT to extract the longest isoform and classify coding and non-coding gene loci. The longest protein sequences were used to improve the reference annotation and perform the orthology analysis described below. We use Infernal v1.1.4 (Nawrocki and Eddy 2013) to find RNA homology from non-coding transcripts in Rfam 14.7 (Kalvari et al. 2017, 2018).

We analyzed nanoPARE reads separately for the analysis of posttranscriptional regulation events. We performed this analysis using annotated miRNAs, 21-nt phasiRNAs, and 24-nt phasiRNAs while assemblies containing the longest transcripts were used to identify sRNA targets. We used PAREsnip2 (Thody et al. 2018) to predict all sRNA-target pairs and to validate the effective sRNA-guided cleavage site with nanoPARE reads. We ran PAREsnip2 withdefault parameters using Fahlgren & Carrington targeting rules (Fahlgren and Carrington 2009). We considered only targets in categories 0 and 1 with P-value < 0.05 present across three replicates for downstream analysis. The protein sequences of target transcripts were analyzed with pfam_scan.pl to search known protein domains. We used the orthologous table to retrieve A. thaliana and O. sativa orthologs. Orthogroups were used to identify targets overlapping between species and we used UpSet plot function from ComplexHeatmap R program v2.18.0 (Gu 2022) to visualize overlap in miRNA and phasiRNA targets in anthers across species. Functional annotation of orthologous genes was used to interpret the role of miRNAs and phasiRNAs in developing anthers.

### Orthology and phylogeny analysis of AGO, DCL and RDR genes

Reference-guided *de novo* transcript assemblies were incorporated into the reference annotations to improve the gene models, complete the gene sets, and annotate AGO, DCL and RDR proteins by inferring phylogenetic trees. To infer accurate phylogenetic trees, and to annotate trees we took advantage of a comprehensive phylogenetic analysis published (Bélanger et al. 2023) by our group using the data from 32 species. The evolutionary relationship between all the examined plants can be visualized in the species tree (Supplemental Figure 6A). We used OrthoFinder v2.5.4 (Emms and Kelly 2015) to perform a gene orthology inference in sampled species. We used the annotation from Bélanger et al. (Bélanger et al. 2023) to identify orthologous groups of AGO, DCL and RDR protein families and performed downstream analyses as described in Bélanger et al. (Bélanger et al. 2023). A total of 667 AGO, 224 DCL and 266 RDR proteins were annotated in the 39 species (Detailed in Supplemental Table 6) and corresponding phylogenetic trees are shown in Supplemental Figure 6 B–D. Of these, we found a total of 255 AGO, 64 DCL and 83 RDR proteins among the sampled species (Supplemental Table 7). To compare and interpret the expression levels of *AGO*, *DCL* and *RDR* genes between developmental stages of anthers, we added the Z-score values of expressed genes to the phylogenetic trees using iTOL (Letunic and Bork 2019)and focused on genes with shared expression patterns across species (Figure 3 and Supplemental Figure 7).

## Data Availability

The complete set of raw sRNA-seq, RNA-seq and nanoPARE reads were deposited in the Sequence Read Archive under SRA accession number PRJNA1107926.

## AUTHOR CONTRIBUTIONS

S.B. and B.C.M designed the experiment; S.B. and J.Z. constructed RNA libraries and performed the experiments; S.B. and Y.Y. analyzed the data; S.B., Y.Y. and B.C.M interpreted results. All authors contributed to writing the manuscript.

## FUNDING INFORMATION

This work was supported by the National Key R&D Program of China (award 2023ZD04073, to J.Z.), USDA, National Institute of Food and Agriculture (“BTT EAGER” award no. 2018–09058 to B.C.M.), as well as resources from the Donald Danforth Plant Science Center and the University of Missouri–Columbia.

## ACKNOWLEDGMENTS

We thank Lynn G. Clark (Iowa State University) for providing bamboo plants. We thank members of the Meyers lab for helpful discussions and Joanna Friesner (Donald Danforth Plant Science Center) for assistance with editing. We thank Mayumi Nakano (Donald Danforth Plant Science Center) for assistance with data handling. S. Bélanger gratefully acknowledges graduate studentships from the National Sciences and Engineering Research Council of Canada (NSERC) and the NSERC CREATE AgroPhytoSciences program.

## COMPETING INTEREST STATEMENT

The authors declare no conflict of interest.

## Notes

### Competing Interest Statement

The authors have declared no competing interest.

### Summary of Updates

The title and abstract were modified.

