## Supplemental Figure 1 for "Comparative RNA profiling identifies stage-specific phasiRNAs and co-expressed *Argonaute* genes in Bambusoideae and Pooideae species"

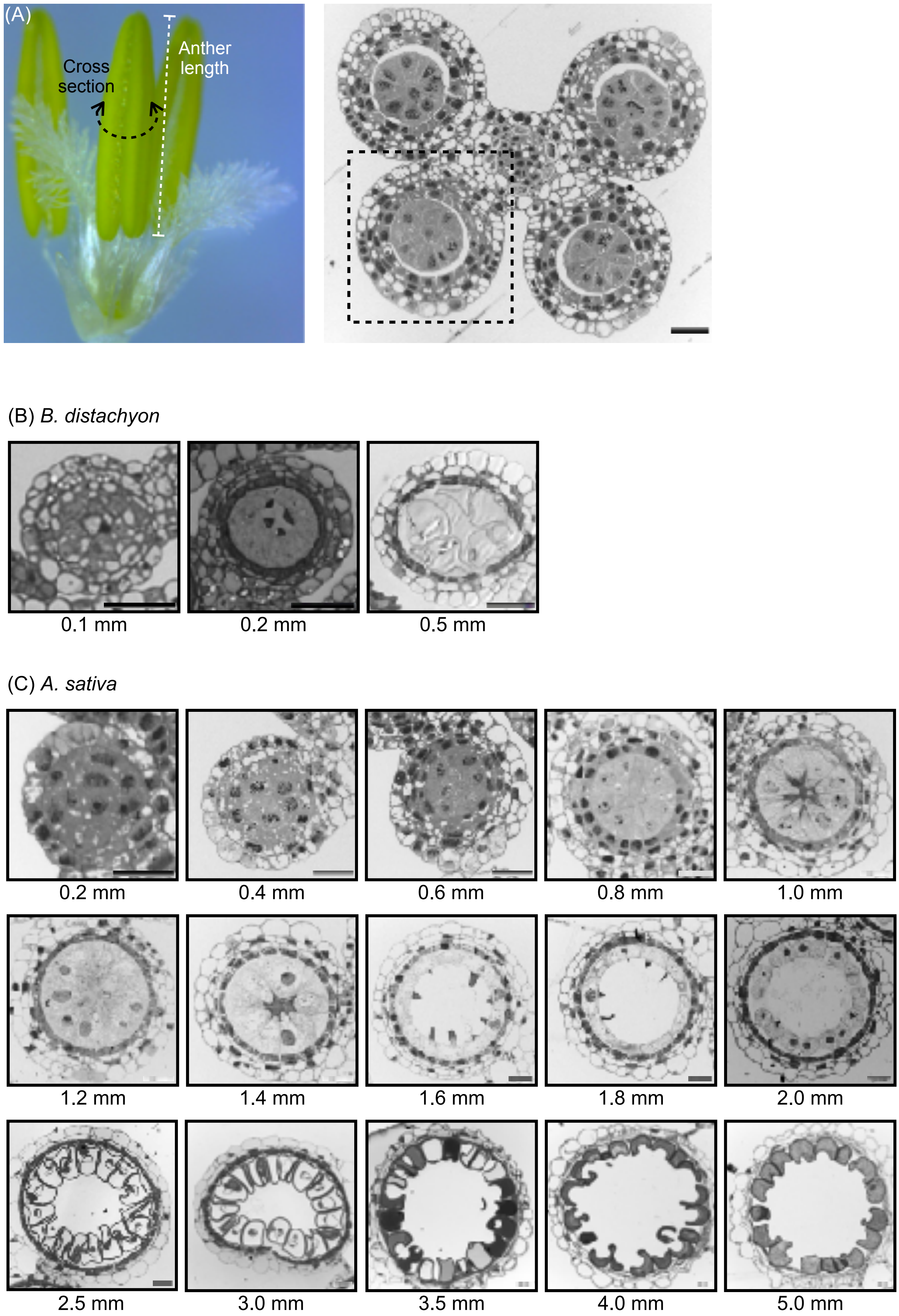
(A)
Cross section
Anther length
(B) B. distachyon
0.1 mm
0.2 mm
0.5 mm
(C) A. sativa
0.2 mm
0.4 mm
0.6 mm
0.8 mm
1.0 mm
1.2 mm
1.4 mm
1.6 mm
1.8 mm
2.0 mm
2.5 mm
3.0 mm
3.5 mm
4.0 mm
5.0 mm

(D) *H. vulgare*

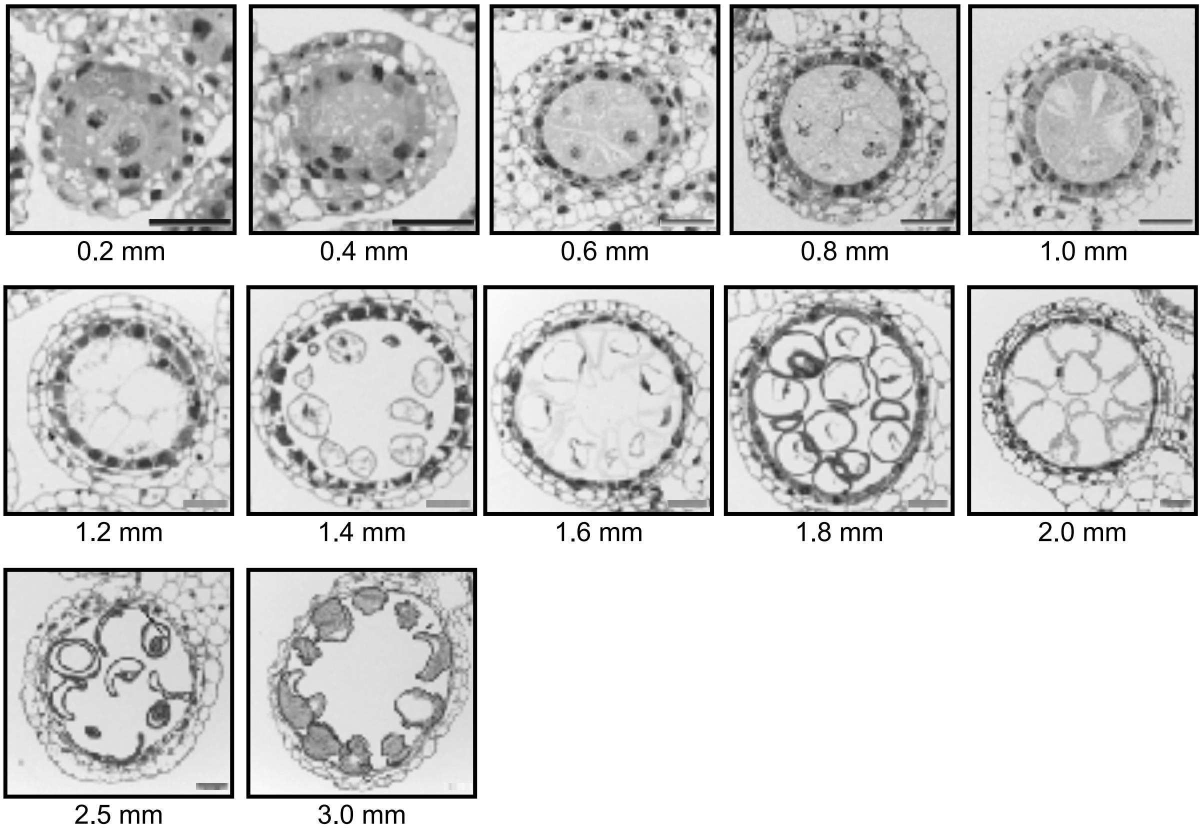


(E) *S. cereale*

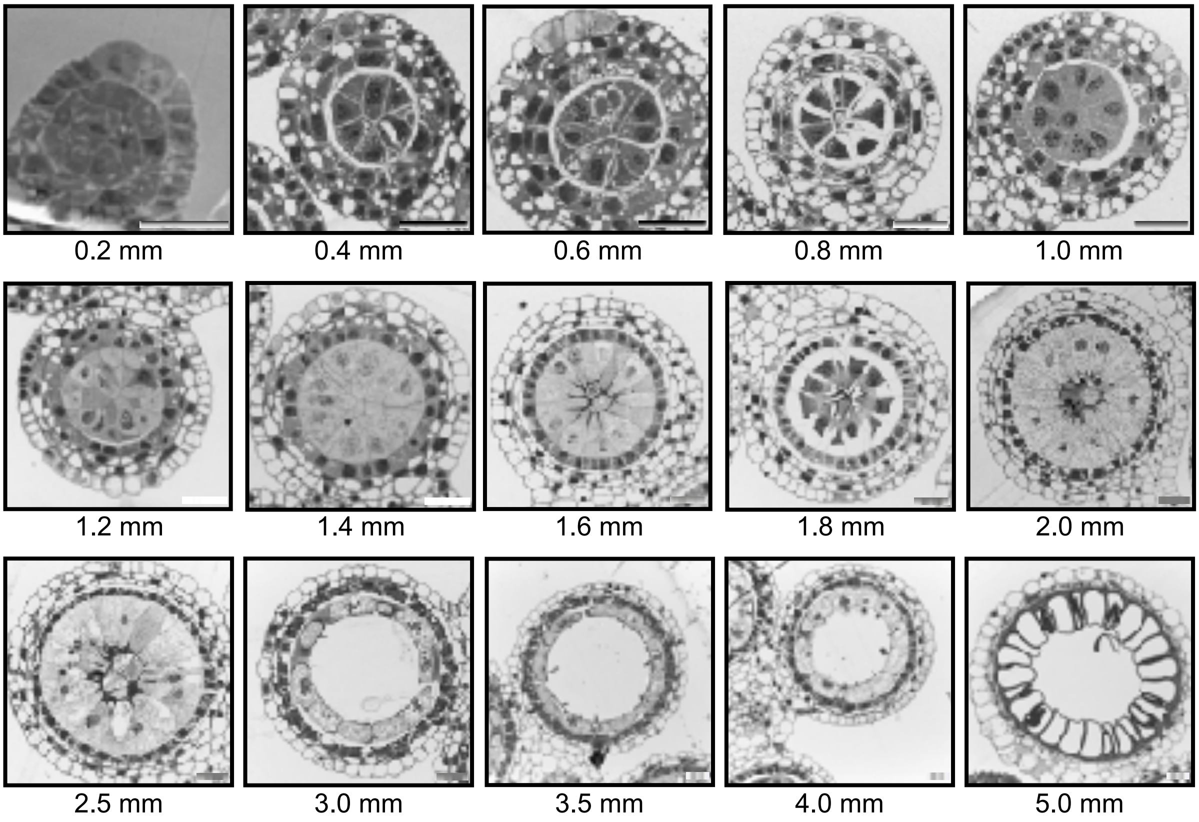

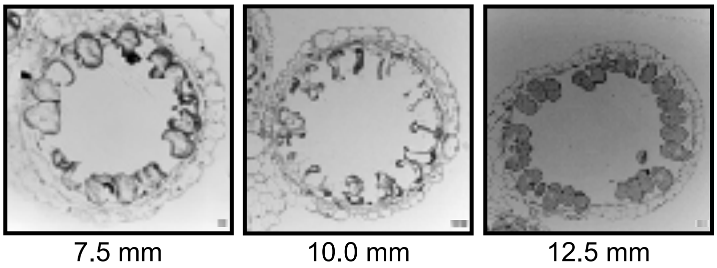
7.5 mm
10.0 mm
12.5 mm

(F) *T. turgidum*

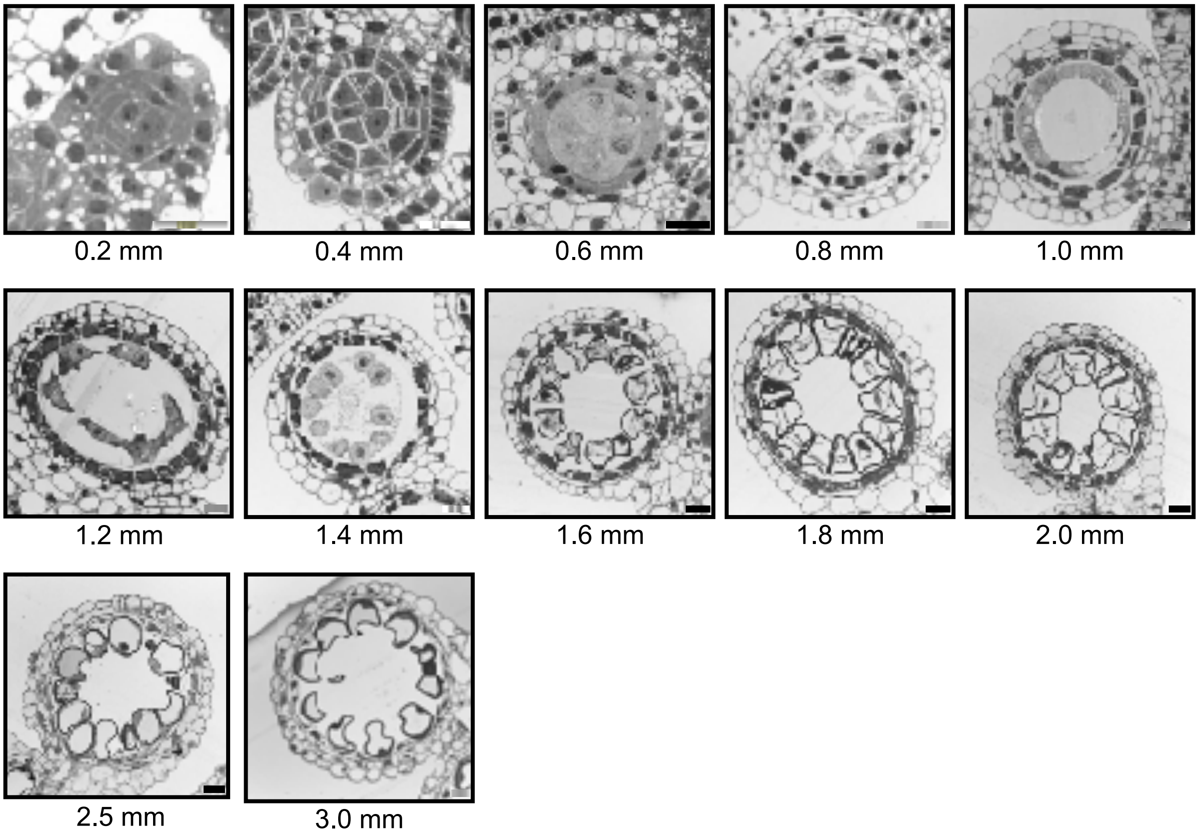
0.2 mm
0.4 mm
0.6 mm
0.8 mm
1.0 mm
1.2 mm
1.4 mm
1.6 mm
1.8 mm
2.0 mm
2.5 mm
3.0 mm

(G) *T. aestivum*

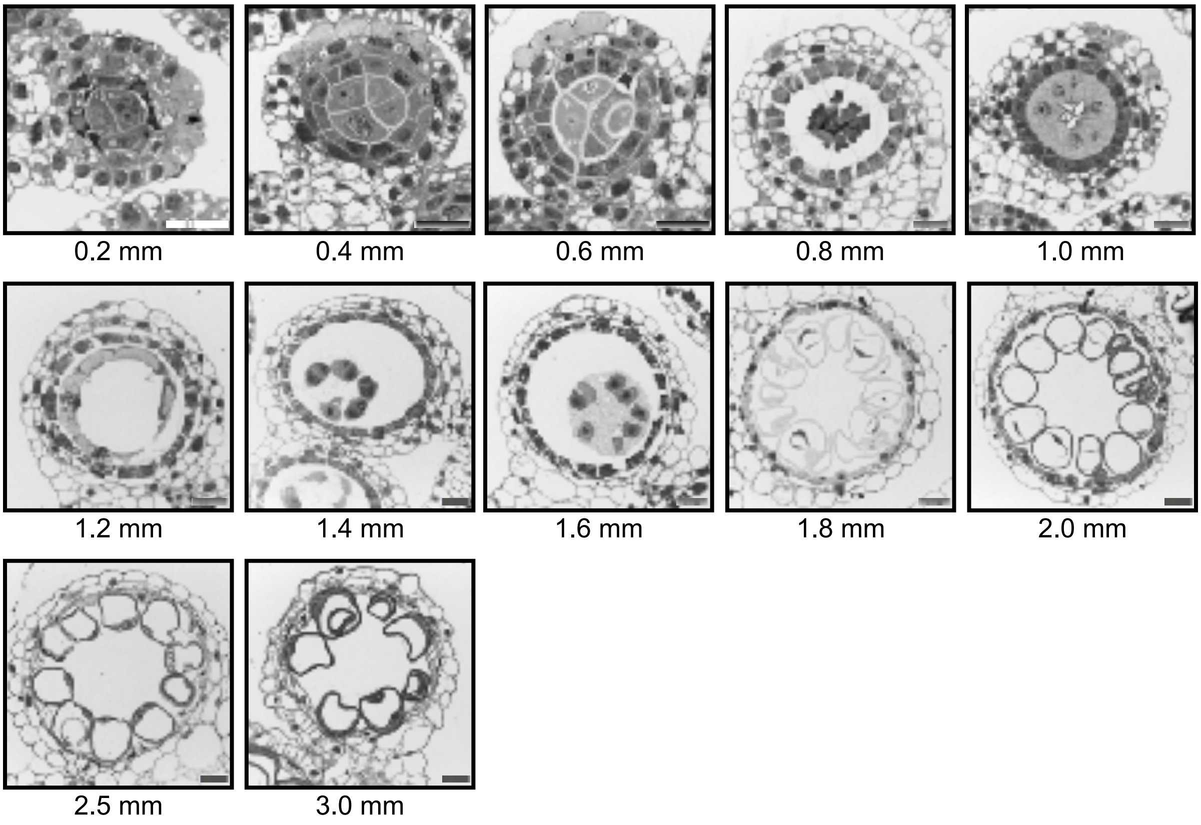

*Supplemental Figure S1. Histological examination of anther development in Pooideae species.*

(A) Cross-sections of anthers were taken from the middle portion of each anther (indicated by black on the left), and images were captured from one lobe of the anther, as shown on the right. Anther length was determined as indicated on the left. Detailed anther sections of (B) *Brachypodium distachyon*, (C) *Avena sativa*, (D) *Hordeum vulgare*, (E) *Secale cereale*, (F) *Triticum turgidum*, and (G) *Triticum aestivum* anthers. Anthers were fixed using a 2% paraformaldehyde:glutaraldehyde solution, embedded in Quetol resin, sectioned to 500 μm, and stained with toluidine blue in epoxy resin. Scale bars are indicated as 20 μm.
