## Supplemental Figure 2 for "Comparative RNA profiling identifies stage-specific phasiRNAs and co-expressed *Argonaute* genes in Bambusoideae and Pooideae species"

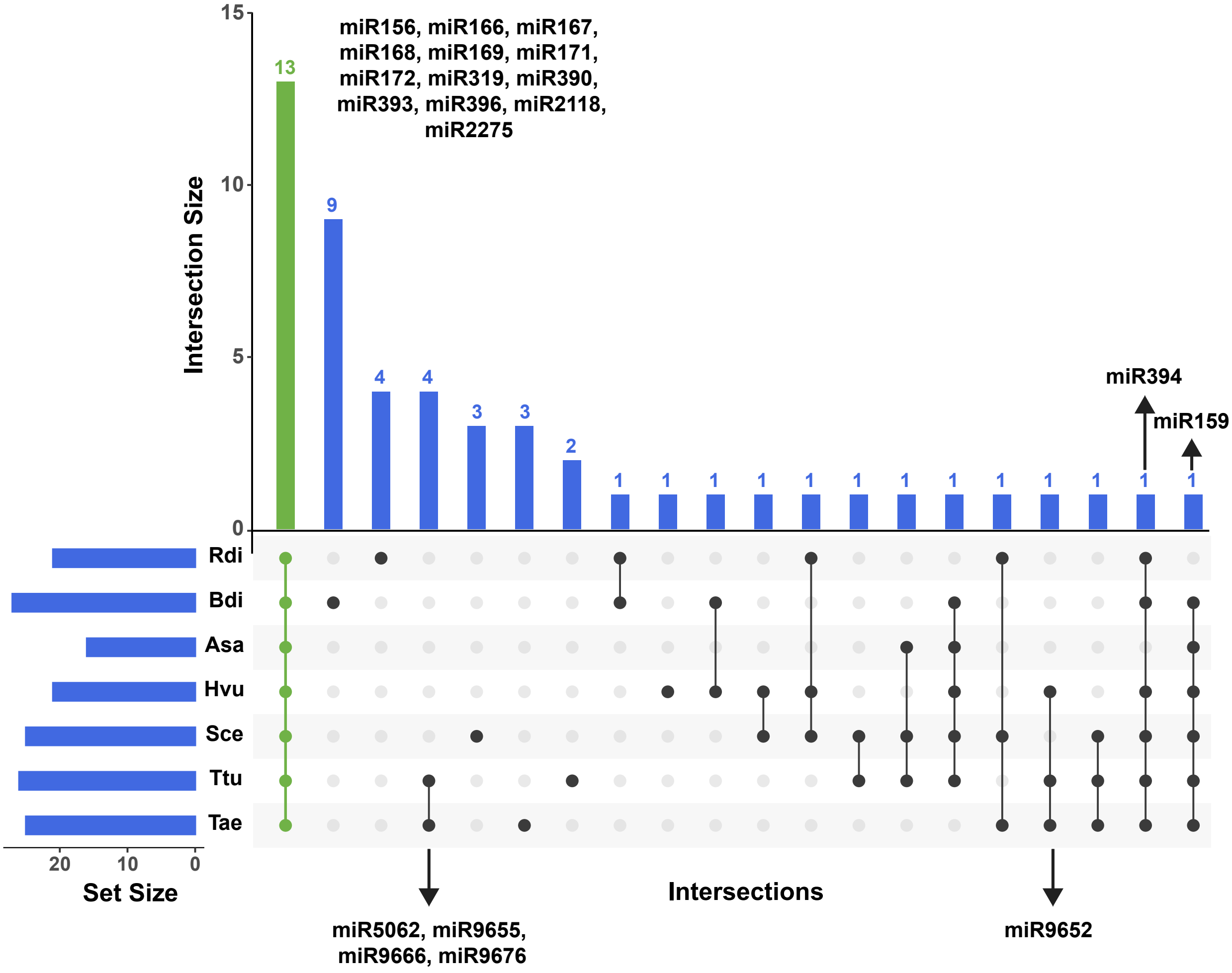

Intersection Size
15
10
5
0
13
9
4
4
3
3
2
1
1
1
1
1
1
1
1
1
1
1
1
1
miR156, miR166, miR167, miR168, miR169, miR171, miR172, miR319, miR390, miR393, miR396, miR2118, miR2275
miR394
miR159
Rdi
Bdi
Asa
Hvu
Sce
Ttu
Tae
20
10
0
Set Size
Intersections
miR5062, miR9655, miR9666, miR9676
miR9652

*Supplemental Figure S2. MicroRNA predictions across sampled species reveal a limited overlap in conserved expression patterns of miRNA families in the anther.*

Upset plots illustrate miRBase annotated miRNAs shared across species. The bottom-left plot presents the size of each set as a horizontal histogram, the bottom-right displays the intersection matrix, and the upper-right indicates the size of each combination as a vertical histogram.
