## Supplemental Figure 3 for "Comparative RNA profiling identifies stage-specific phasiRNAs and co-expressed *Argonaute* genes in Bambusoideae and Pooideae species"

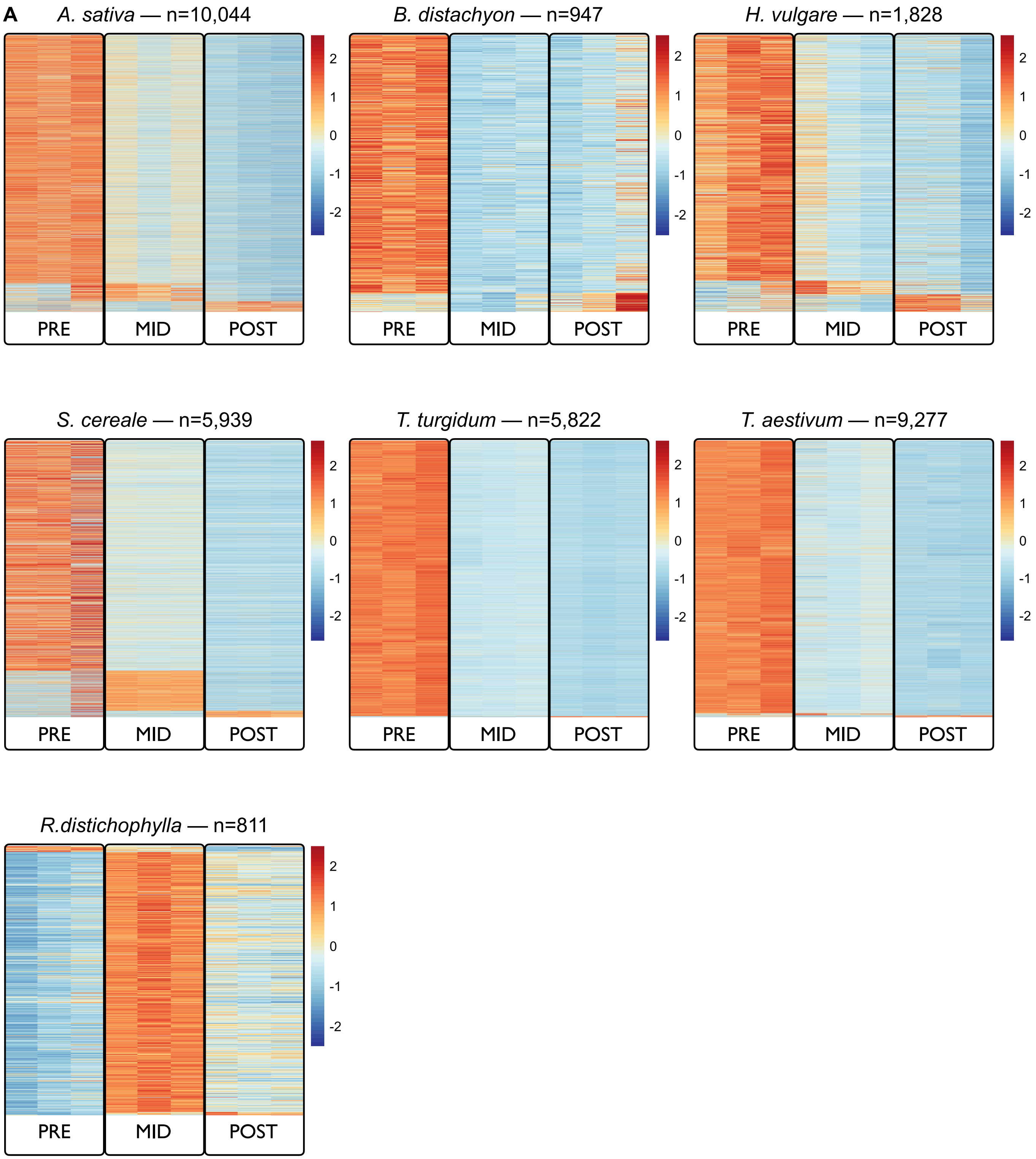
A
A. sativa — n=10,044
B. distachyon — n=947
H. vulgare — n=1,828
S. cereale — n=5,939
T. turgidum — n=5,822
T. aestivum — n=9,277
R.distichophylla — n=811
PRE
MID
POST
2
1
0
-1
-2

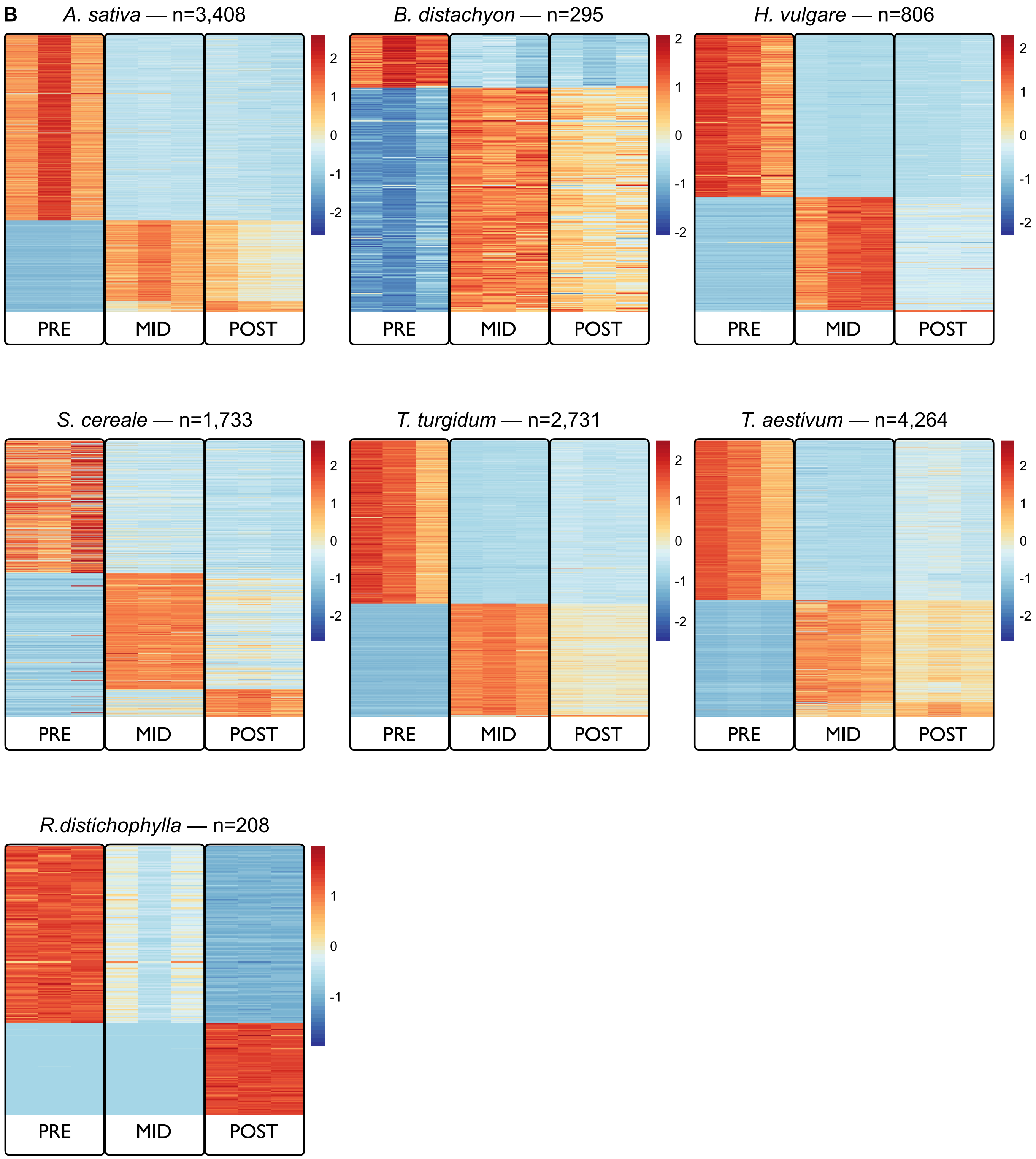
B
A. sativa — n=3,408
B. distachyon — n=295
H. vulgare — n=806
S. cereale — n=1,733
T. turgidum — n=2,731
T. aestivum — n=4,264
R.distichophylla — n=208
PRE
MID
POST
2
1
0
-1
-2

*Supplemental Figure S3. Accumulation of 21-nt (A) and 24-nt (B) phasiRNAs annotated in the anthers of seven species.*

The heatmap illustrates the accumulation pattern of phasiRNAs, with the scale bar indicating the relative abundance change between columns, representing a relative change.
