## Supplemental Figure 4 for "Comparative RNA profiling identifies stage-specific phasiRNAs and co-expressed *Argonaute* genes in Bambusoideae and Pooideae species"

*R. distichophylla*
Length: 22; E-value: 1.1e-054; n= 52/137

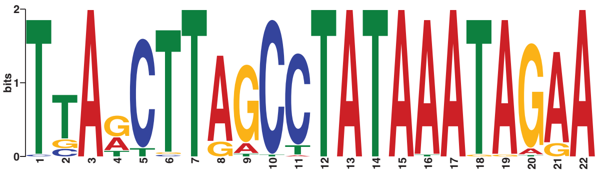

*B. distachyon*
Length: 21; E-value: 2.2e-028; n= 43/56

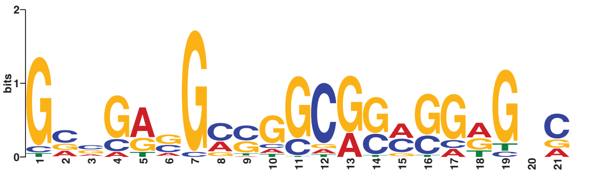

*A. sativa*
Length: 22; E-value: 5.1e-211; n= 1900/2284

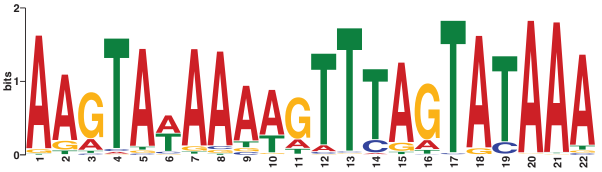

*H. vulgare*
Length: 22; E-value: 1.1e-155; n= 388/472

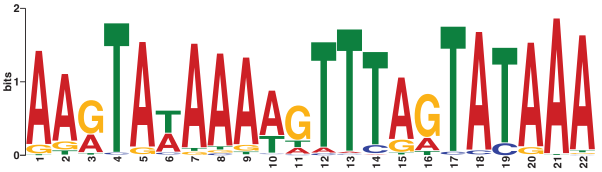

*S. cereale*
Length: 22; E-value: 5.4e-180; n= 501/829

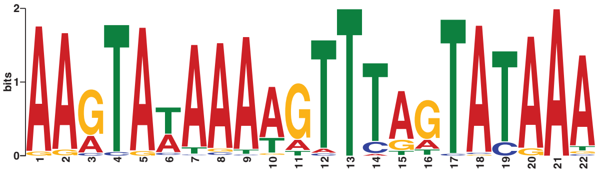

*T. turgidum subsp durum*
Length: 22; E-value: 3.2e-214; n= 1365/1610

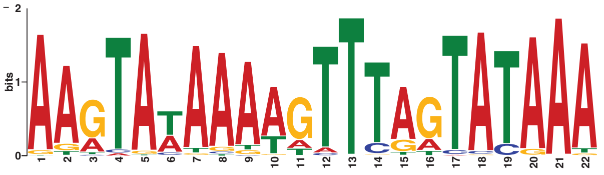

*T. aestivum*
Length: 22; E-value: 4.0e-235; n= 1323/2457

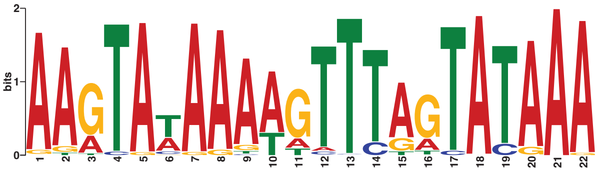

*R. distichophylla*
Length: 22; E-value: 5.8e-199; n= 61/71

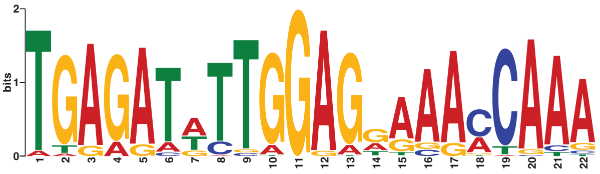

*B. distachyon*
Length: 22; E-value: 5.7e-246; n= 217/239

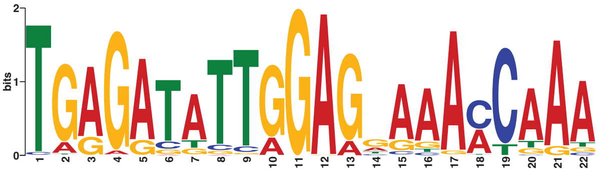

*A. sativa*
Length: 22; E-value: 2.5e-231; n= 1095/1124

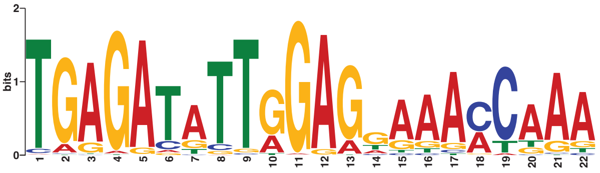

*H. vulgare*
Length: 22; E-value: 1.6e-185; n= 298/334

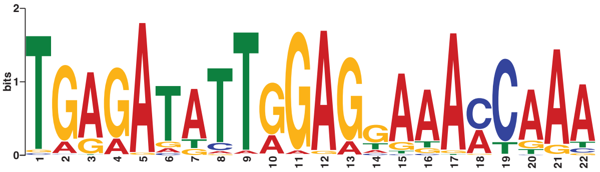

*S. cereale*
Length: 22; E-value: 2.2e-199; n= 647/904

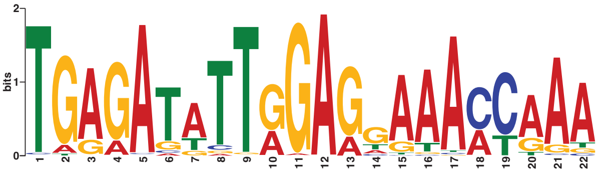

*T. turgidum subsp durum*
Length: 22; E-value: 1.5e-221; n= 919/1121

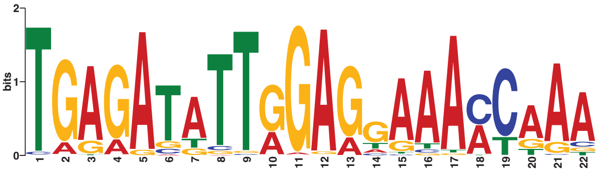

*T. aestivum*
Length: 22; E-value: 1.6e-194; n= 1485/1807

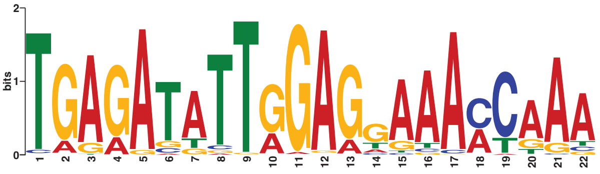

*Supplemental Figure S4. Conserved motifs found in putative 24-*PHAS *transcripts for premeiotic (left; unknown motif) and meiotic (right; miR2275 motif) phasiRNAs across seven species.*

On the right panel the motif found in *R. distichophylla* corresponds to postmeiotic 24-*PHAS* loci.
