## Supplemental Figure 5 for "Comparative RNA profiling identifies stage-specific phasiRNAs and co-expressed *Argonaute* genes in Bambusoideae and Pooideae species"

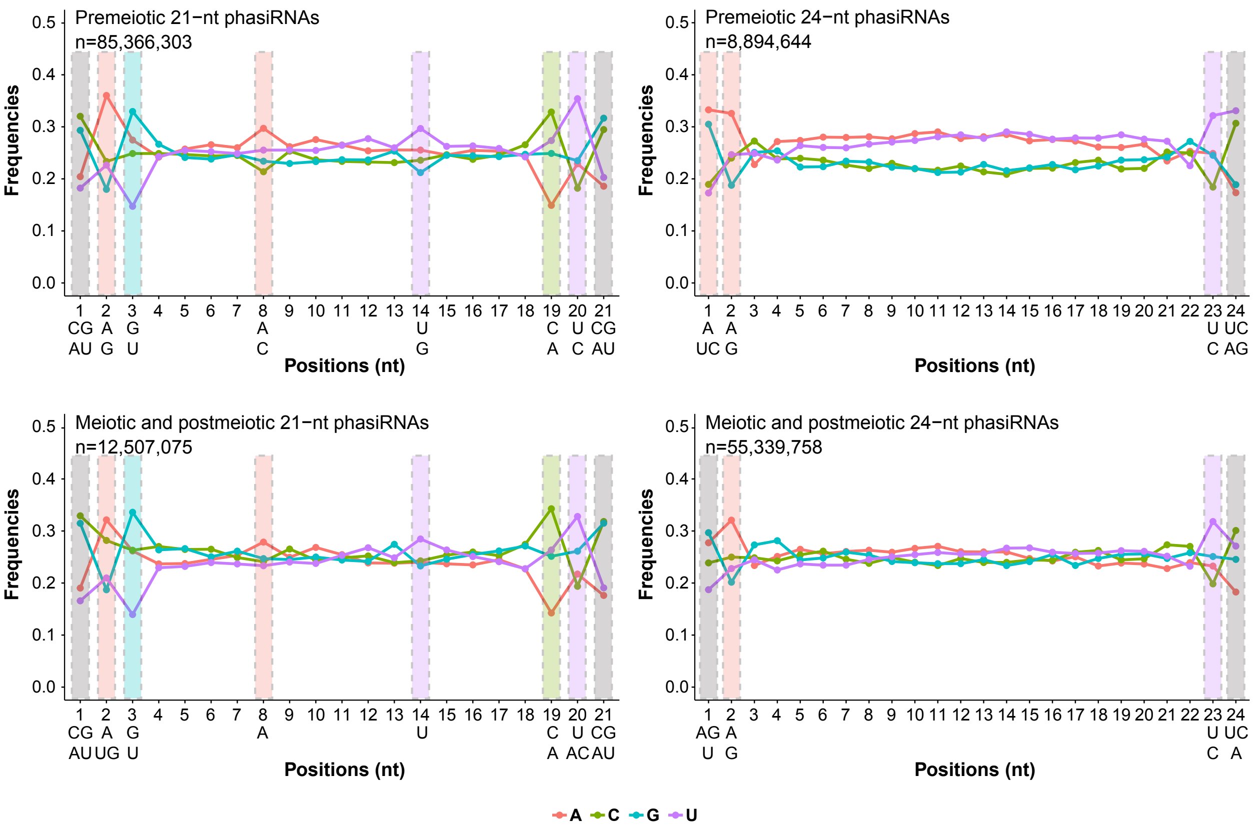

Premeiotic 21−nt phasiRNAs
n=85,366,303
Premeiotic 24−nt phasiRNAs
n=8,894,644
Meiotic and postmeiotic 21−nt phasiRNAs
n=12,507,075
Meiotic and postmeiotic 24−nt phasiRNAs
n=55,339,758
Frequencies
Positions (nt)
A
C
G
U

*Supplemental Figure S5. Positional and compositional nucleotide signatures observed in reproductive phasiRNAs.*

We computed the nucleotide frequency at each position along 21-nt (left) and 24-nt (right) phasiRNAs. The frequency was calculated using all phasiRNAs expressed in *PHAS* loci across all species. The positional and compositional nucleotide signatures are conserved between groups of premeiotic (upper) and meiotic/postmeiotic (lower) in both 21-nt and 24-nt phasiRNAs. Specific nucleotides are conserved at the 3' end positions of both 21-nt and 24-nt phasiRNAs, but not at the 5' end. The y-axis represents the frequency of nucleotides distributed at each position of 21-nt and 24-nt phasiRNAs (x-axis). Nucleotides selected for further discussion in the main text are highlighted in colored rectangles.
