## Supplemental Figure 6 for "Comparative RNA profiling identifies stage-specific phasiRNAs and co-expressed *Argonaute* genes in Bambusoideae and Pooideae species"

A

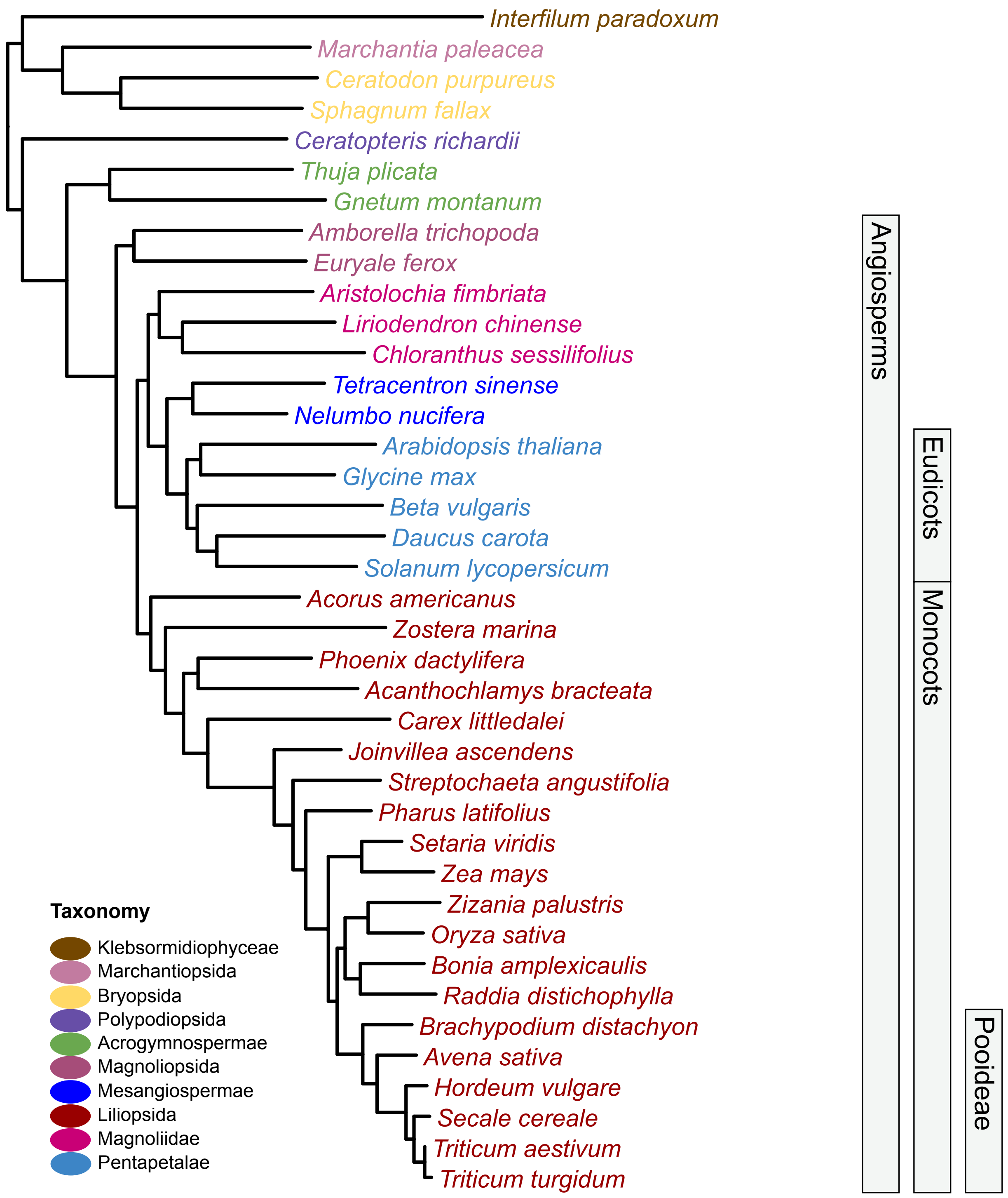
Interfilum paradoxum
Marchantia paleacea
Ceratodon purpureus
Sphagnum fallax
Ceratopteris richardii
Thuja plicata
Gnetum montanum
Amborella trichopoda
Euryale ferox
Aristolochia fimbriata
Liriodendron chinense
Chloranthus sessilifolius
Tetracentron sinense
Nelumbo nucifera
Arabidopsis thaliana
Glycine max
Beta vulgaris
Daucus carota
Solanum lycopersicum
Acorus americanus
Zostera marina
Phoenix dactylifera
Acanthochlamys bracteata
Carex littledalei
Joinvillea ascendens
Streptochaeta angustifolia
Pharus latifolius
Setaria viridis
Zea mays
Zizania palustris
Oryza sativa
Bonia amplexicaulis
Raddia distichophylla
Brachypodium distachyon
Avena sativa
Hordeum vulgare
Secale cereale
Triticum aestivum
Triticum turgidum
Angiosperms
Eudicots
Monocots
Pooideae
Taxonomy
Klebsormidiophyceae
Marchantiopsida
Bryopsida
Polypodiopsida
Acrogymnospermae
Magnoliopsida
Mesangiospermae
Liliopsida
Magnoliidae
Pentapetalae

B

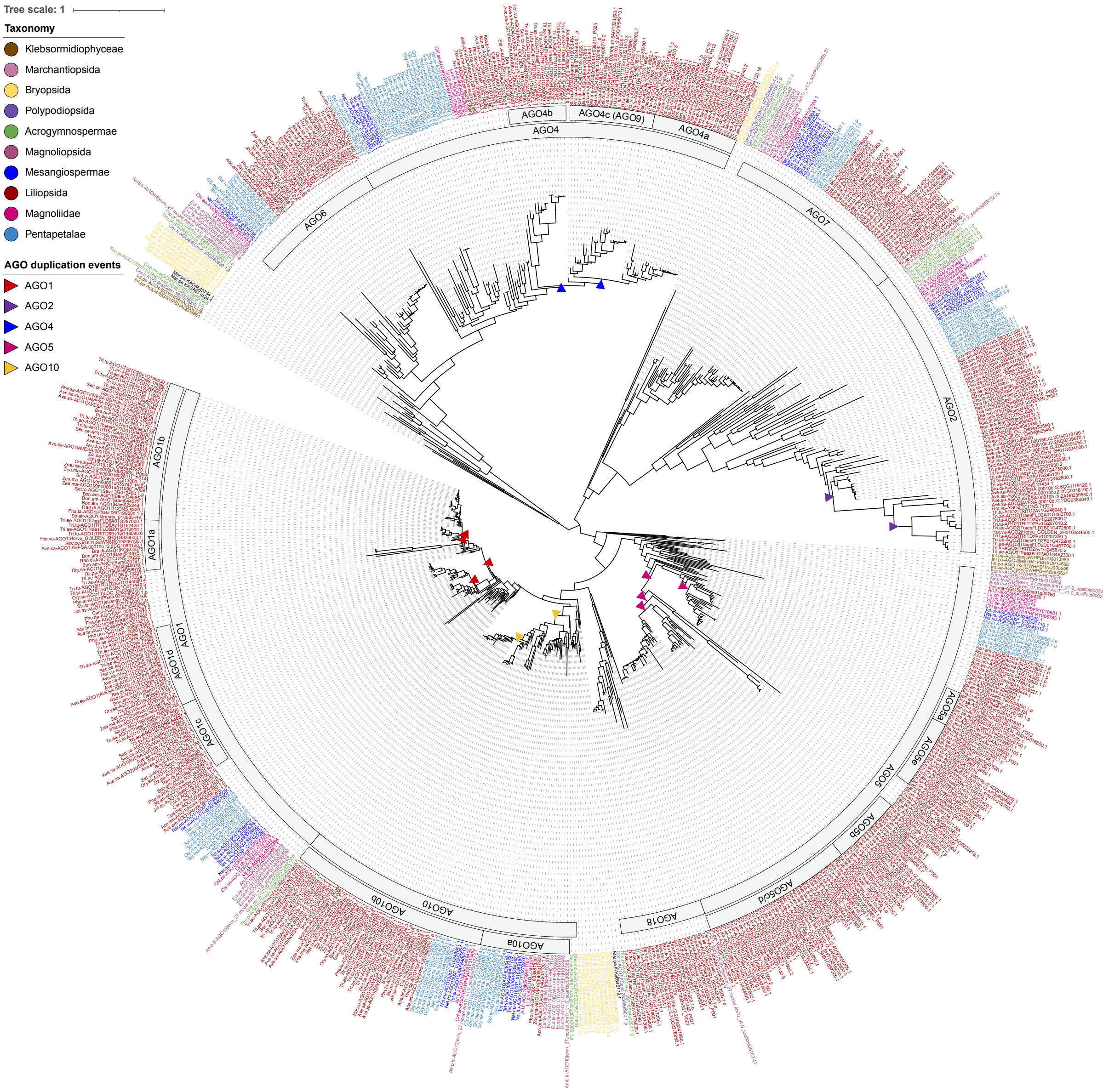


667 AGO proteins

**C**

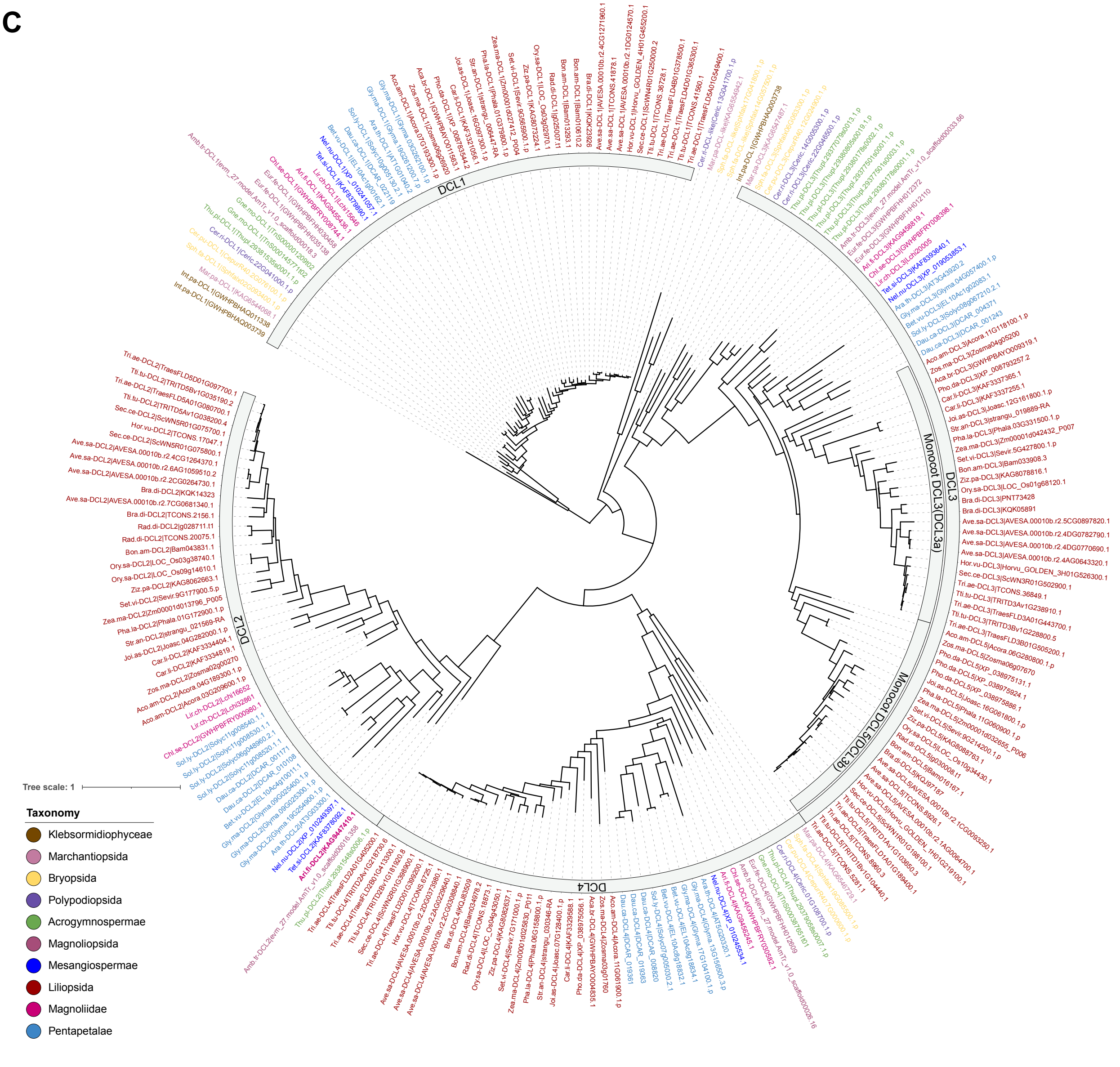


**224 DCL proteins**

**D**

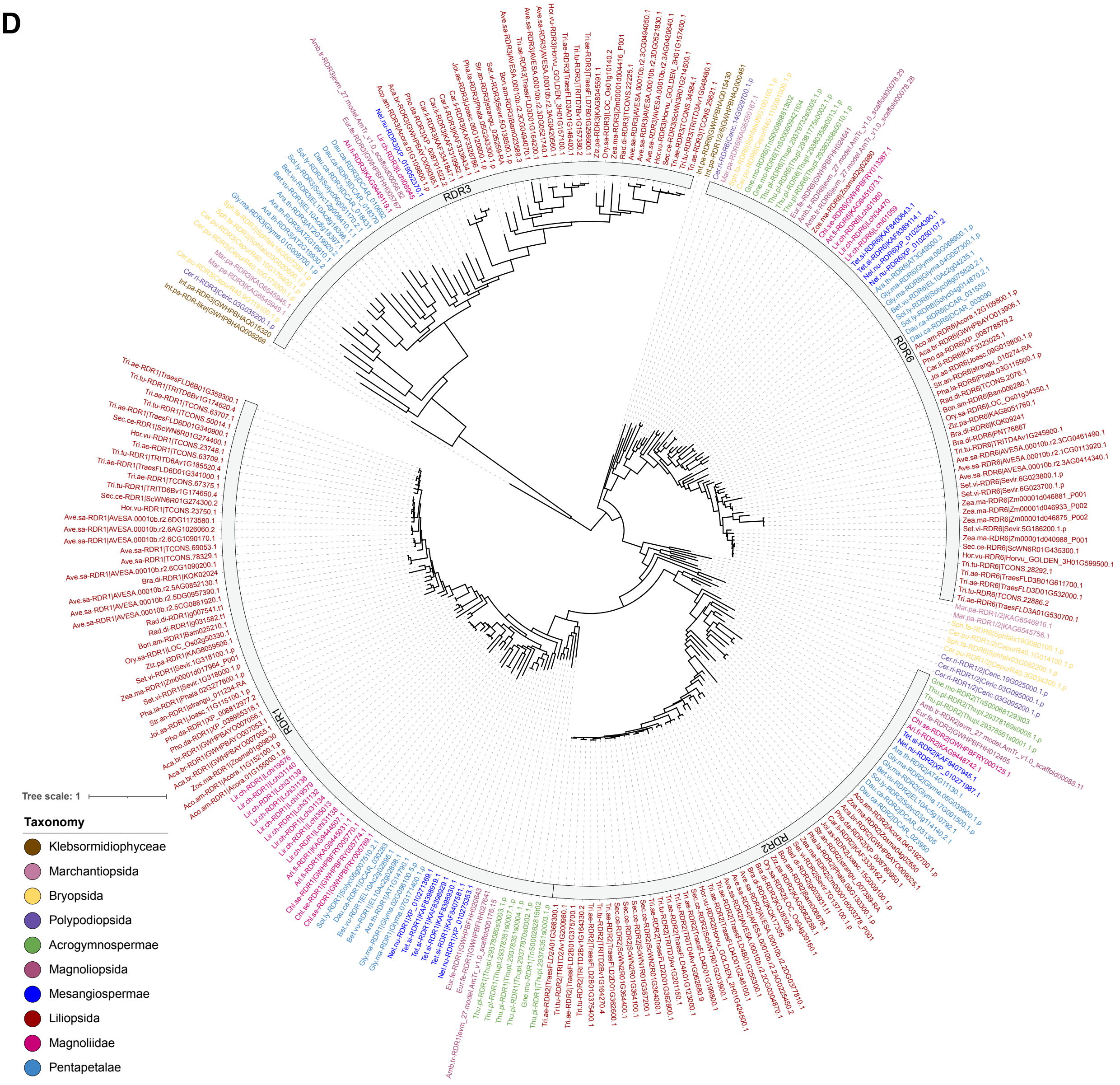


**266 RDR proteins**

*Supplemental Figure S6. Phylogenetic trees inferred from 39 species.*

(A) Species tree of all the examined species in this study. The tree was generated using OrthoFinder based on whole proteome sequences and visualized using iTOL. (B-D) Phylogenetic trees inferred for AGO, DCL and RDR proteins annotated among the 39 species
