## Supplemental Figure 7 for "Comparative RNA profiling identifies stage-specific phasiRNAs and co-expressed *Argonaute* genes in Bambusoideae and Pooideae species"

**A**

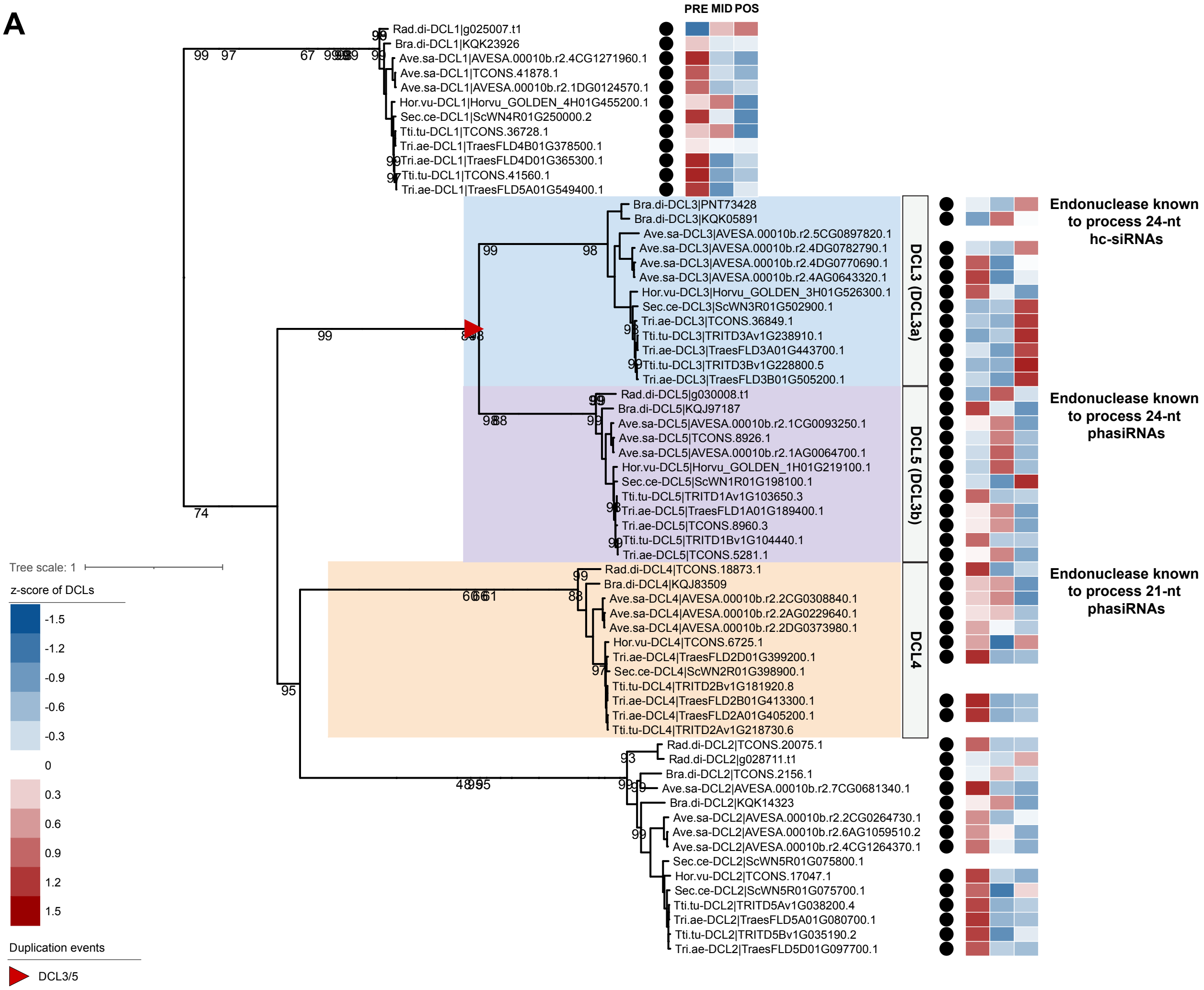
PRE MID POS
Rad.di-DCL1|g025007.t1
Bra.di-DCL1|KQK23926
Ave.sa-DCL1|AVESA.00010b.r2.4CG1271960.1
Ave.sa-DCL1|TCONS.41878.1
Ave.sa-DCL1|AVESA.00010b.r2.1DG0124570.1
Hor.vu-DCL1|Horvu_GOLDEN_4H01G455200.1
Sec.ce-DCL1|ScWN4R01G250000.2
Tti.tu-DCL1|TCONS.36728.1
Tri.ae-DCL1|TraesFLD4B01G378500.1
Tri.ae-DCL1|TraesFLD4D01G365300.1
Tti.tu-DCL1|TCONS.41560.1
Tri.ae-DCL1|TraesFLD5A01G549400.1
Bra.di-DCL3|PNT73428
Bra.di-DCL3|KQK05891
Ave.sa-DCL3|AVESA.00010b.r2.5CG0897820.1
Ave.sa-DCL3|AVESA.00010b.r2.4DG0782790.1
Ave.sa-DCL3|AVESA.00010b.r2.4DG0770690.1
Ave.sa-DCL3|AVESA.00010b.r2.4AG0643320.1
Hor.vu-DCL3|Horvu_GOLDEN_3H01G526300.1
Sec.ce-DCL3|ScWN3R01G502900.1
Tri.ae-DCL3|TCONS.36849.1
Tti.tu-DCL3|TRITD3Av1G238910.1
Tri.ae-DCL3|TraesFLD3A01G443700.1
Tti.tu-DCL3|TRITD3Bv1G228800.5
Tri.ae-DCL3|TraesFLD3B01G505200.1
Rad.di-DCL5|g030008.t1
Bra.di-DCL5|KQJ97187
Ave.sa-DCL5|AVESA.00010b.r2.1CG0093250.1
Ave.sa-DCL5|TCONS.8926.1
Ave.sa-DCL5|AVESA.00010b.r2.1AG0064700.1
Hor.vu-DCL5|Horvu_GOLDEN_1H01G219100.1
Sec.ce-DCL5|ScWN1R01G198100.1
Tti.tu-DCL5|TRITD1Av1G103650.3
Tri.ae-DCL5|TraesFLD1A01G189400.1
Tri.ae-DCL5|TCONS.8960.3
Tti.tu-DCL5|TRITD1Bv1G104440.1
Tri.ae-DCL5|TCONS.5281.1
Rad.di-DCL4|TCONS.18873.1
Bra.di-DCL4|KQJ83509
Ave.sa-DCL4|AVESA.00010b.r2.2CG0308840.1
Ave.sa-DCL4|AVESA.00010b.r2.2AG0229640.1
Ave.sa-DCL4|AVESA.00010b.r2.2DG0373980.1
Hor.vu-DCL4|TCONS.6725.1
Tri.ae-DCL4|TraesFLD2D01G399200.1
Sec.ce-DCL4|ScWN2R01G398900.1
Tti.tu-DCL4|TRITD2Bv1G181920.8
Tri.ae-DCL4|TraesFLD2B01G413300.1
Tri.ae-DCL4|TraesFLD2A01G405200.1
Tti.tu-DCL4|TRITD2Av1G218730.6
Rad.di-DCL2|TCONS.20075.1
Rad.di-DCL2|g028711.t1
Bra.di-DCL2|TCONS.2156.1
Ave.sa-DCL2|AVESA.00010b.r2.7CG0681340.1
Bra.di-DCL2|KQK14323
Ave.sa-DCL2|AVESA.00010b.r2.2CG0264730.1
Ave.sa-DCL2|AVESA.00010b.r2.6AG1059510.2
Ave.sa-DCL2|AVESA.00010b.r2.4CG1264370.1
Sec.ce-DCL2|ScWN5R01G075800.1
Hor.vu-DCL2|TCONS.17047.1
Sec.ce-DCL2|ScWN5R01G075700.1
Tti.tu-DCL2|TRITD5Av1G038200.4
Tri.ae-DCL2|TraesFLD5A01G080700.1
Tti.tu-DCL2|TRITD5Bv1G035190.2
Tri.ae-DCL2|TraesFLD5D01G097700.1
DCL3 (DCL3a)
DCL5 (DCL3b)
DCL4
Endonuclease known to process 24-nt hc-siRNAs
Endonuclease known to process 24-nt phasiRNAs
Endonuclease known to process 21-nt phasiRNAs
Tree scale: 1
z-score of DCLs
-1.5
-1.2
-0.9
-0.6
-0.3
0
0.3
0.6
0.9
1.2
1.5
Duplication events
DCL3/5

B

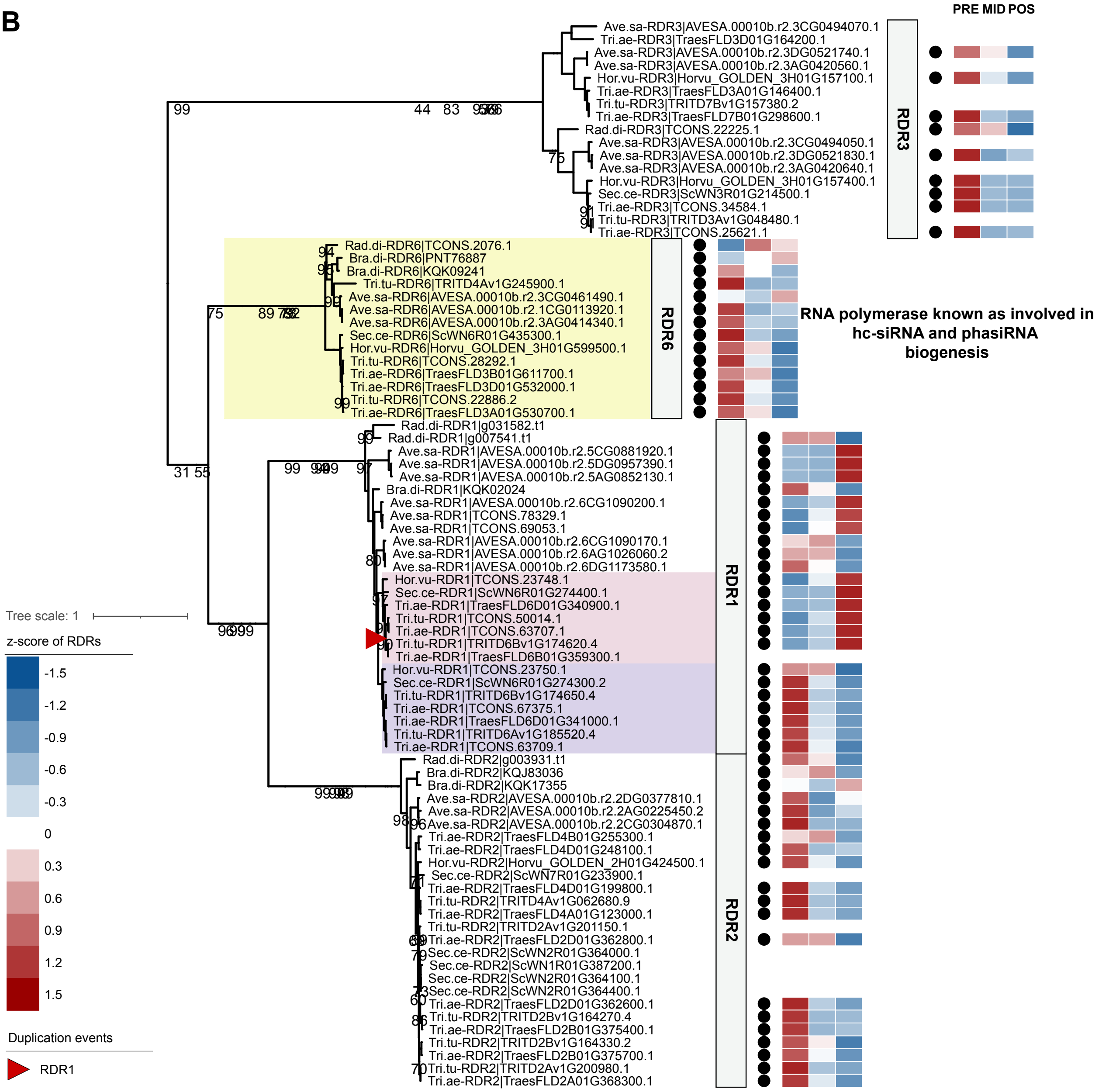

PRE MID POS
Ave.sa-RDR3|AVESA.00010b.r2.3CG0494070.1
Tri.ae-RDR3|TraesFLD3D01G164200.1
Ave.sa-RDR3|AVESA.00010b.r2.3DG0521740.1
Ave.sa-RDR3|AVESA.00010b.r2.3AG0420560.1
Hor.vu-RDR3|Horvu_GOLDEN_3H01G157100.1
Tri.ae-RDR3|TraesFLD3A01G146400.1
Tri.tu-RDR3|TRITD7Bv1G157380.2
Tri.ae-RDR3|TraesFLD7B01G298600.1
Rad.di-RDR3|TCONS.22225.1
Ave.sa-RDR3|AVESA.00010b.r2.3CG0494050.1
Ave.sa-RDR3|AVESA.00010b.r2.3DG0521830.1
Ave.sa-RDR3|AVESA.00010b.r2.3AG0420640.1
Hor.vu-RDR3|Horvu_GOLDEN_3H01G157400.1
Sec.ce-RDR3|ScWN3R01G214500.1
Tri.ae-RDR3|TCONS.34584.1
Tri.tu-RDR3|TRITD3Av1G048480.1
Tri.ae-RDR3|TCONS.25621.1
RDR3
Rad.di-RDR6|TCONS.2076.1
Bra.di-RDR6|PNT76887
Bra.di-RDR6|KQK09241
Tri.tu-RDR6|TRITD4Av1G245900.1
Ave.sa-RDR6|AVESA.00010b.r2.3CG0461490.1
Ave.sa-RDR6|AVESA.00010b.r2.1CG0113920.1
Ave.sa-RDR6|AVESA.00010b.r2.3AG0414340.1
Sec.ce-RDR6|ScWN6R01G435300.1
Hor.vu-RDR6|Horvu_GOLDEN_3H01G599500.1
Tri.tu-RDR6|TCONS.28292.1
Tri.ae-RDR6|TraesFLD3B01G611700.1
Tri.ae-RDR6|TraesFLD3D01G532000.1
Tri.tu-RDR6|TCONS.22886.2
Tri.ae-RDR6|TraesFLD3A01G530700.1
RDR6
RNA polymerase known as involved in hc-siRNA and phasiRNA biogenesis
Rad.di-RDR1|g031582.t1
Rad.di-RDR1|g007541.t1
Ave.sa-RDR1|AVESA.00010b.r2.5CG0881920.1
Ave.sa-RDR1|AVESA.00010b.r2.5DG0957390.1
Ave.sa-RDR1|AVESA.00010b.r2.5AG0852130.1
Bra.di-RDR1|KQK02024
Ave.sa-RDR1|AVESA.00010b.r2.6CG1090200.1
Ave.sa-RDR1|TCONS.78329.1
Ave.sa-RDR1|TCONS.69053.1
Ave.sa-RDR1|AVESA.00010b.r2.6CG1090170.1
Ave.sa-RDR1|AVESA.00010b.r2.6AG1026060.2
Ave.sa-RDR1|AVESA.00010b.r2.6DG1173580.1
Hor.vu-RDR1|TCONS.23748.1
Sec.ce-RDR1|ScWN6R01G274400.1
Tri.ae-RDR1|TraesFLD6D01G340900.1
Tri.tu-RDR1|TCONS.50014.1
Tri.ae-RDR1|TCONS.63707.1
Tri.tu-RDR1|TRITD6Bv1G174620.4
Tri.ae-RDR1|TraesFLD6B01G359300.1
Hor.vu-RDR1|TCONS.23750.1
Sec.ce-RDR1|ScWN6R01G274300.2
Tri.tu-RDR1|TRITD6Bv1G174650.4
Tri.ae-RDR1|TCONS.67375.1
Tri.ae-RDR1|TraesFLD6D01G341000.1
Tri.tu-RDR1|TRITD6Av1G185520.4
Tri.ae-RDR1|TCONS.63709.1
RDR1
Rad.di-RDR2|g003931.t1
Bra.di-RDR2|KQJ83036
Bra.di-RDR2|KQK17355
Ave.sa-RDR2|AVESA.00010b.r2.2DG0377810.1
Ave.sa-RDR2|AVESA.00010b.r2.2AG0225450.2
Ave.sa-RDR2|AVESA.00010b.r2.2CG0304870.1
Tri.ae-RDR2|TraesFLD4B01G255300.1
Tri.ae-RDR2|TraesFLD4D01G248100.1
Hor.vu-RDR2|Horvu_GOLDEN_2H01G424500.1
Sec.ce-RDR2|ScWN7R01G233900.1
Tri.ae-RDR2|TraesFLD4D01G199800.1
Tri.tu-RDR2|TRITD4Av1G062680.9
Tri.ae-RDR2|TraesFLD4A01G123000.1
Tri.tu-RDR2|TRITD2Av1G201150.1
Tri.ae-RDR2|TraesFLD2D01G362800.1
Sec.ce-RDR2|ScWN2R01G364000.1
Sec.ce-RDR2|ScWN1R01G387200.1
Sec.ce-RDR2|ScWN2R01G364100.1
Sec.ce-RDR2|ScWN2R01G364400.1
Tri.ae-RDR2|TraesFLD2D01G362600.1
Tri.tu-RDR2|TRITD2Bv1G164270.4
Tri.ae-RDR2|TraesFLD2B01G375400.1
Tri.tu-RDR2|TRITD2Bv1G164330.2
Tri.ae-RDR2|TraesFLD2B01G375700.1
Tri.tu-RDR2|TRITD2Av1G200980.1
Tri.ae-RDR2|TraesFLD2A01G368300.1
RDR2
Tree scale: 1
z-score of RDRs
-1.5
-1.2
-0.9
-0.6
-0.3
0
0.3
0.6
0.9
1.2
1.5
Duplication events
RDR1

*Supplemental Figure S7. The phylogenetic tree and expression profiles of genes encoding DCL (A) and RDR (B) proteins in the sampled species.*

The phylogenetic tree was derived from the maximum-likelihood analysis based on 39 species. Clades representing DCL and RDR proteins are clearly indicated. Genes expressing DCL and RDR proteins in anthers are marked with a black dot and the heatmap denoting the calculated z-score relative expression change between premeiotic (inside), meiotic (middle) and postmeiotic (outside) stages. Protein clades with known or predicted functions in phasiRNA biogenesis and function are also highlighted.
