## Supplemental Figure 8 for "Comparative RNA profiling identifies stage-specific phasiRNAs and co-expressed *Argonaute* genes in Bambusoideae and Pooideae species"

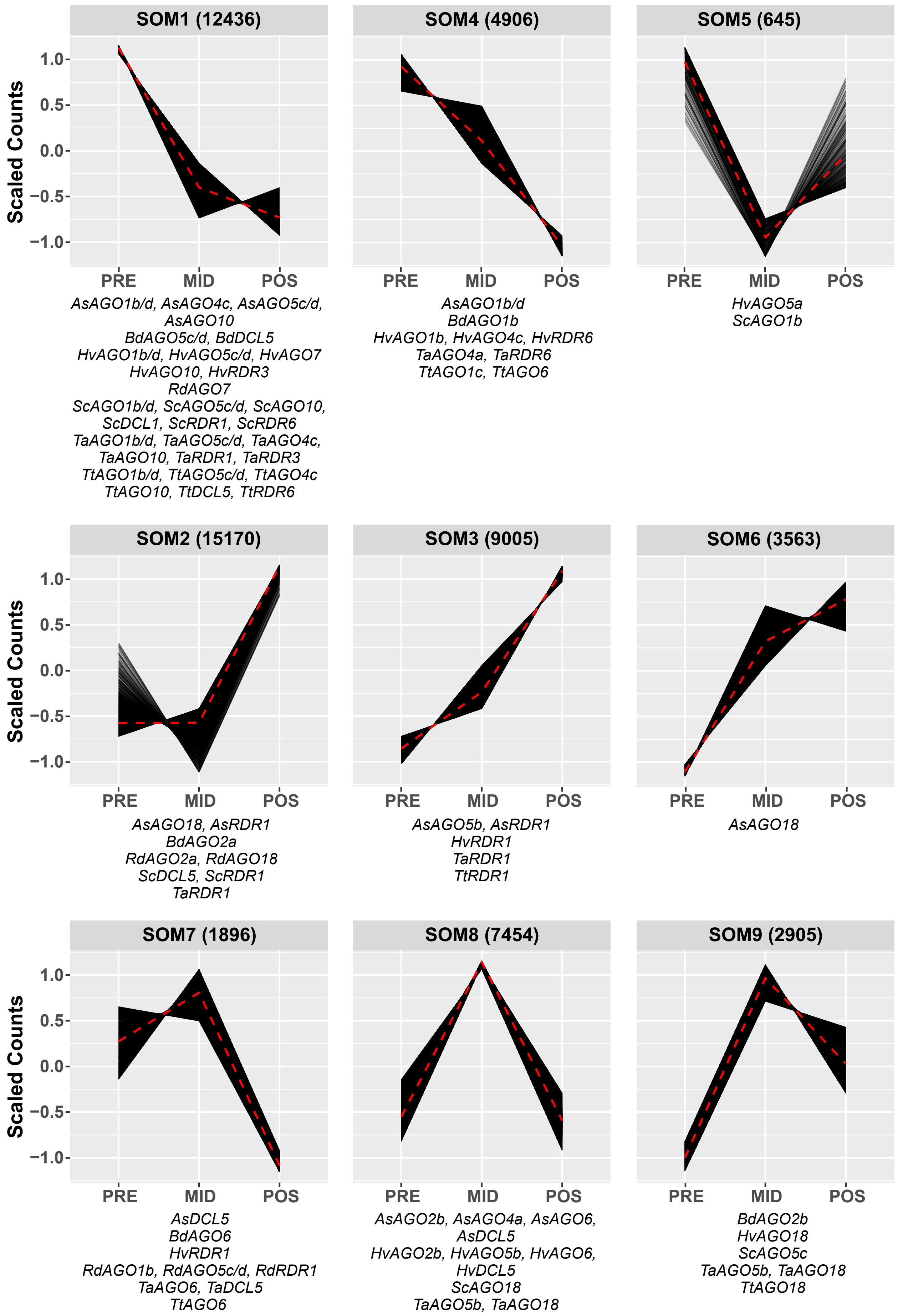
SOM1 (12436)
SOM4 (4906)
SOM5 (645)
Scaled Counts
1.0
0.5
0.0
−0.5
−1.0
PRE
MID
POS
*AsAGO1b/d, AsAGO4c, AsAGO5c/d, AsAGO10*
*BdAGO5c/d, BdDCL5*
*HvAGO1b/d, HvAGO5c/d, HvAGO7*
*HvAGO10, HvRDR3*
*RdAGO7*
*ScAGO1b/d, ScAGO5c/d, ScAGO10,*
*ScDCL1, ScRDR1, ScRDR6*
*TaAGO1b/d, TaAGO5c/d, TaAGO4c,*
*TaAGO10, TaRDR1, TaRDR3*
*TtAGO1b/d, TtAGO5c/d, TtAGO4c*
*TtAGO10, TtDCL5, TtRDR6*
*AsAGO1b/d*
*BdAGO1b*
*HvAGO1b, HvAGO4c, HvRDR6*
*TaAGO4a, TaRDR6*
*TtAGO1c, TtAGO6*
*HvAGO5a*
*ScAGO1b*
SOM2 (15170)
SOM3 (9005)
SOM6 (3563)
*AsAGO18, AsRDR1*
*BdAGO2a*
*RdAGO2a, RdAGO18*
*ScDCL5, ScRDR1*
*TaRDR1*
*AsAGO5b, AsRDR1*
*HvRDR1*
*TaRDR1*
*TtRDR1*
*AsAGO18*
SOM7 (1896)
SOM8 (7454)
SOM9 (2905)
*AsDCL5*
*BdAGO6*
*HvRDR1*
*RdAGO1b, RdAGO5c/d, RdRDR1*
*TaAGO6, TaDCL5*
*TtAGO6*
*AsAGO2b, AsAGO4a, AsAGO6,*
*AsDCL5*
*HvAGO2b, HvAGO5b, HvAGO6,*
*HvDCL5*
*ScAGO18*
*TaAGO5b, TaAGO18*
*BdAGO2b*
*HvAGO18*
*ScAGO5c*
*TaAGO5b, TaAGO18*
*TtAGO18*

*Supplemental Figure S8. Coexpressed genes in the anthers of seven species.*

The clustering analysis reveals nine clusters, displaying the relative abundance of differentially expressed gene modules, alongside annotations for AGO, DCL, and RDR genes within those modules. Differential expression and gene clustering analyses were conducted using the R packages DESeq2 and KOHONEN, respectively.
